# Genome structure of *Brachionus asplanchnoidis*, a Eukaryote with intrapopulation variation in genome size

**DOI:** 10.1101/2021.03.09.434534

**Authors:** C.P. Stelzer, J. Blommaert, A.M. Waldvogel, M. Pichler, B. Hecox-Lea, D.B. Mark Welch

## Abstract

Eukaryotic genomes vary greatly in size due to variation in the proportion of non-coding DNA, a pattern that emerges both in comparisons at a larger taxonomic scale and at the level of individuals within a species. The rotifer *Brachionus asplanchnoidis* represents one of the most extreme cases of intraspecific genome size variation among Eukaryotes, displaying almost 2-fold variation within a geographic population. Here we used a whole-genome sequencing approach to identify the underlying DNA sequence differences by assembling a high-quality reference genome draft for one individual of the population and aligning short-reads of 15 individuals from the same geographic population. We identified large, contiguous copy number variable regions (CNVs), which exhibited significant coverage differences among individuals, and whose coverage overall scaled with genome size. CNVs were mainly composed of tandemly repeated satellite DNA, with only few interspersed genes or other sequences, and were characterized by an elevated GC-content. Judging from their distributions across contigs, some CNVs are fragments of accessory (B-)chromosomes while others resemble large insertions to normal chromosomes. CNV patterns in offspring of two parents with divergent genome size, and CNV patterns in several individuals from an inbred line differing in genome size demonstrated inheritance and accumulation of CNVs across generations. Our study provides unprecedented insights into genome size evolution at microevolutionary time scales and thus paves the way for studying genome size evolution in contemporary populations rather than inferring patterns and processes *a posteriori* from species comparisons.

## Introduction

The genomes of Eukaryotic organisms display remarkable diversity in size, overall spanning approximately five orders of magnitude (Elliott and Gregory 2015). In addition, genome size may vary substantially among closely related species (Stelzer et al. 2011; Jeffery et al. 2016), within a species (e.g., Šmarda et al. 2008; Ruiz-Ruano et al. 2011; Chia et al. 2012), and sometimes even within a population (Stelzer et al. 2019). Most of the variation in genome size stems from differences in the proportion of various kinds of non-coding DNA and/or transposable elements, which can reach excessive levels in species with giant genomes (Shah et al. 2020; Meyer et al. 2021). Studying genome size variation at the DNA-sequence level allows identification of exactly those genomic elements that make up for the genome size difference, and it can suggest the relative strength of mutation, selection, and drift - the underlying evolutionary forces ultimately causing divergence in genome size.

Much of our understanding of eukaryotic genome size variation comes from comparisons between closely related species. Recent studies suggested that proliferation of repetitive elements (RE), in particular transposable elements, play an important role in genome expansion, while their silencing and deletion has been implicated in the streamlining of genomes. In a few studies, it was possible to pinpoint individual REs as the driver of genome expansion (Naville et al. 2019; Wong et al. 2019), whereas in other studies, differently-sized genomes were found to differ in several classes of REs (Blommaert et al. 2019; McCann et al. 2020). In the latter case, it is difficult to decide whether multiple RE classes have expanded more or less simultaneously in evolutionary time, or whether the expansions of some REs have occurred *after* an initial genome size divergence driven by a single element. Without accurate dating of the expansions of individual elements, interspecific comparisons suffer from such a ‘blind spot’ on the early stages of genome divergence. Ultimately, all genome size differences must have gone through a stage of intrapopulation variation followed by fixation, or loss, of these size-variants. Thus, identifying genome size variants within populations and studying them on microevolutionary time scales may allow additional insights on the evolutionary dynamics of early genome divergences.

Intraspecific genome size variation (IGV) has been described in several species of Eukaryotes (some examples are summarized in Smarda and Bures 2010; Stelzer et al. 2019). IGV may be associated with variation in the number of chromosomes (e.g., B-chromosomes Smarda and Bures 2010), but there are also examples where IGV is not reflected in the karyotype (Šmarda et al. 2008; Jeffery et al. 2016). One of the most recent additions to IGV model organisms was the monogonont rotifer *B. asplanchnoidis*, which displays a nearly 2-fold variation in genome size even among individuals within a geographic population (Stelzer et al. 2019). Monogonont rotifers are short-lived (1-2 weeks), small aquatic metazoans, only a few hundred micrometers in size, common in fresh and brackish water habitats throughout the world. They have a life-cycle involving cyclical parthenogenesis (Nogrady et al. 1993) reproducing by ameiotic parthenogenesis for prolonged periods and inducing sexual reproduction occasionally. A ‘rotifer clone’ consists of the asexual descendants of a single female that has hatched from a single resting egg, which itself is the product of sexual reproduction. In many Monogonont species, sex is triggered by crowding due to accumulation of mixis-inducing peptides released by the animals (Gilbert 2017). In lab cultures, it is possible to suppress sexual reproduction by frequent dilution intervals or large culture volumes, and to induce mixis in small culture volumes or through the use of media drawn from dense cultures. Thus, it is possible to either deliberately cross two rotifer clones sexually, or to keep them clonally for hundreds of generations.

In the present study, we focus on a population of *B. asplanchnoidis* from Obere Halbjochlacke (OHJ), a shallow alkaline lake in Eastern Austria (Riss et al. 2017; Stelzer et al. 2019). Individuals of this population can be crossed with each other - even if they substantially differ in genome size - and they will produce offspring with intermediate genome sizes close to the parental mean. Genome size can be artificially selected up or down with a heritability of 1 by breeding only individuals with large or small genome size. Genome size variation in this system is mediated by relatively large genomic elements (several megabases in size), which segregate independently from each other during meiosis. The smallest observed genome size in *B. asplanchnoidis* was 404Mb (2C, nuclear DNA content). Individuals at or close to this basal genome size are completely lacking independently segregating elements, while in larger individuals, genome size scales with the amount of independently segregating elements (Stelzer et al. 2019).

Here we used a whole-genome sequencing approach to identify the DNA sequence differences responsible for intrapopulation genome size variation in this population of *B. asplanchnoidis*. Our specific goal was to identify and characterize genomic regions that are present in one or multiple copies in some individuals of the OHJ-population, but are missing in others. To this end, we assembled a highly contiguous draft genome of a reference clone using long-read (PacBio) technology and then mapped short-reads of 15 different clones with genome sizes from 404 to 644 Mbp to this assembly. To identify copy number variations (CNVs), we scanned for regions of increased per-bp read coverage. To independently confirm CNVs, we used PCR to detect presence/absence of selected CNVs across different clones of the OHJ-population, and droplet digital PCR to determine the exact copy numbers of one specific locus. Finally, we annotated genes and repetitive elements in the reference genome, and compared CNV regions to non-variable regions of the genome.

## Results

### De novo assembly and annotation of the reference genome

The rotifer clone (OHJ7i3n10) chosen for our reference genome derives from the natural isolate OHJ7 after three rounds of selfing (i.e., fertilizing sexual females by males of the same clone). As measured by flow cytometry (Stelzer et al. 2019), OHJ7i3n10 has a 2C-genome size of 568 Mbp, and thus contains approximately 40% excess DNA, compared to the smallest genome size of the OHJ-population (∼410 Mbp). The total length of our reference assembly was 230.12 Mb, with 455 contigs and an N50 value of 3.065 Mb (**Fig. 1, Supplemental file 1**). Average GC content was 30.5%. However, the GC distribution was not unimodal, but showed two major peaks at ∼25% and ∼35% GC, and a minor one at ∼50% GC (**Supplemental Fig. S1**).

**Fig. 1.**
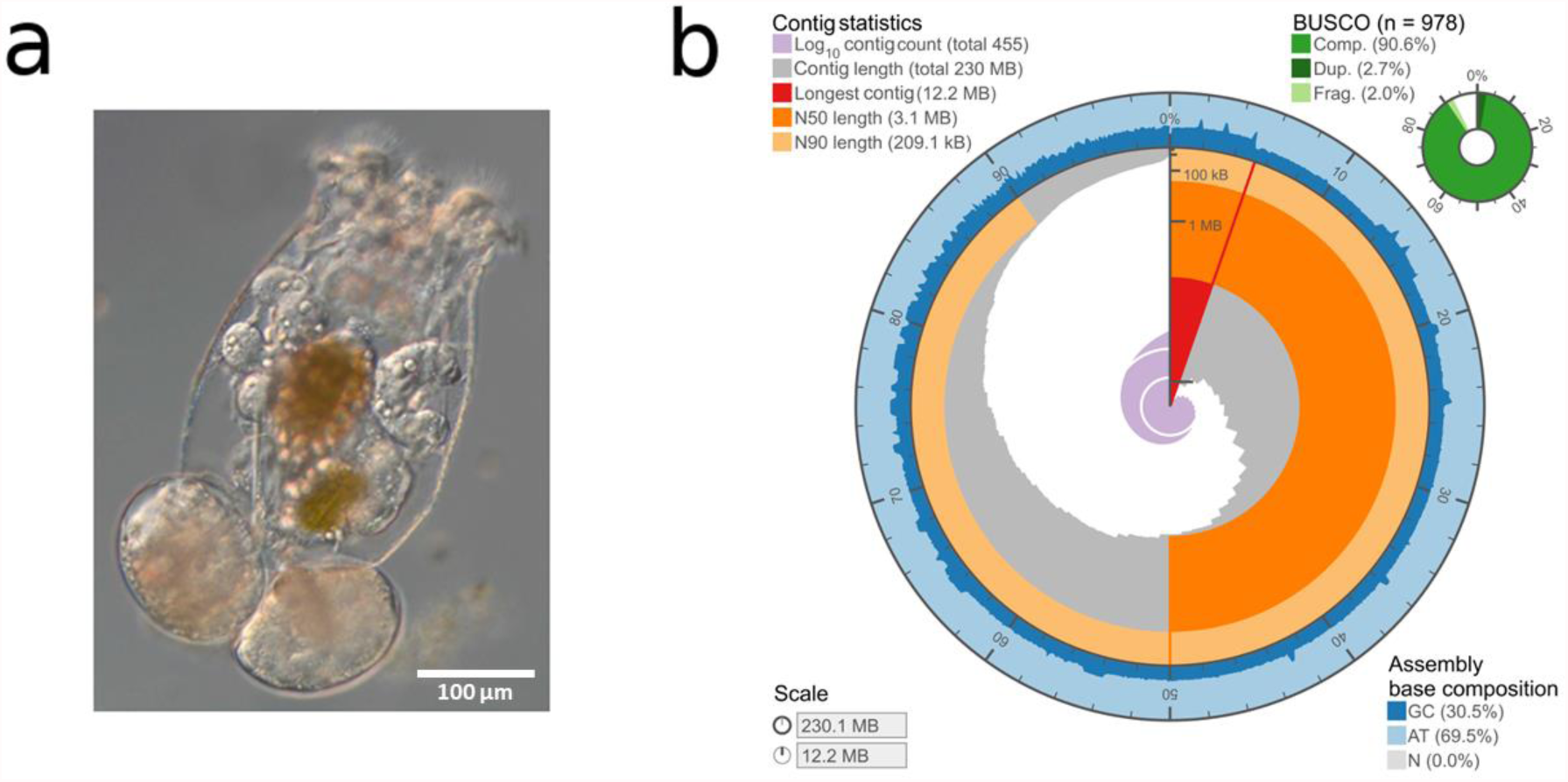
**a** Photograph of *B. asplanchnoidis* female (with two attached asexual eggs). **b** Circular visualization of the contiguity and completeness of the genome assembly. Full circle represents the full assembly length of 230.1 Mb with the longest contig of 12.2 Mb in red and very few contigs <100 kB. GC-content is varying along the assembly (blue).

Taxonomic partitioning of the polished genome assembly confirmed its purity. Most hits could correctly be assigned to rotifers and remaining hits assigned to mollusks and arthropods can mostly be explained by imbalanced availability of rotifer entries in the *nt* database (**Supplemental Fig. S2, Supplemental file 1**). We observed that 90.6 % of the metazoan BUSCO gene set collection was complete with low levels of duplicated (2.7 %), fragmented (2.0 %), and missing (7.4 %) genes (**Supplemental Tab. S2**). A visualization of assembly contiguity and completeness was generated via assembly-stats (Challis 2017) and is presented in **Fig. 1**. Protein-coding genes make up approx. 26 % of the genome assembly length. In total, we annotated 16,667 genes with a median gene length of 1999 bp and approx. five exons per gene (**Supplemental Tab. S3**).

### Comparison of short-reads in 15 OHJ clones

To examine within-population genome size variation, we sequenced 29 short-read libraries from 15 different rotifer clones from the OHJ population (1–4 libraries per clone using different methods, described below). Nine clones were asexual descendants of individuals collected from the field, four (including the source of the reference genome) were each asexual descendants of the same three rounds of selfing of one of these clones, one was an asexual descendant from three rounds of selfing of a different clone, and one was an asexual descendant of a cross between clones derived from crossing two different selfed lineages from two natural isolates (Tab. 1, Tab. S4). Raw reads were passed through multiple preprocessing steps that included quality trimming, removal of PCR duplicates and mitochondrial DNA, and removal of contaminant DNA. Overall, preprocessing reduced the total sequence amount from 265.6 to 194.3 Gbp, resulting in per-base sequencing coverage of 9.4 to 79-fold for the different libraries, with the majority of libraries being above 20-fold (**Supplemental note, Supplemental Fig. S3, Supplemental file 2**).

**Table 1.** Rotifer clones used in this study.

| Clone | Genome Size<sup>1</sup><br>(Mbp) | Origin | No. of<br>libraries |
| --- | --- | --- | --- |
| OHJ82 | 404 | natural clone | 2 |
| OHJ22 | 412 | natural clone | 2 |
| OHJ104 | 462 | natural clone | 2 |
| OHJ97 | 470 | natural clone | 2 |
| OHJ96 | 492 | natural clone | 1 |
| OHJ98 | 504 | natural clone | 1 |
| OHJ105 | 520 | natural clone | 2 |
| OHJ7 | 532 | natural clone | 4 |
| OHJ13 | 536 | natural clone | 2 |
| OHJ22i3n14 | 420 | clone derived from selfing<sup>3</sup> OHJ22 | 1 |
| OHJ7i3n7 | 536 | clone derived from selfing<sup>3</sup> OHJ7 | 2 |
| OHJ7i3n2 | 560 | clone derived from selfing<sup>3</sup> OHJ7 | 2 |
| OHJ7i3n10<sup>2</sup> | 568 | clone derived from selfing<sup>3</sup> OHJ7 | 2 |
| OHJ7i3n5 | 644 | clone derived from selfing<sup>3</sup> OHJ7 | 2 |
| IK1 | 500 | clone from cross of OHJ7i3n2 and OHJ22i3n14 | 2 |

<sup>1</sup> 2C genome size estimated by flow cytometry (Stelzer et al. 2019)
<sup>2</sup> same clone as reference genome
<sup>3</sup> hatched from an individual resting egg after three generations of selfing

Pooling short reads from all libraries revealed the same three-peak pattern of GC-content apparent in the reference assembly, with GC maxima at 26%, 36%, and 48% (**Fig. 2a**). Since these three GC peaks are indicative of three discrete fractions among the genomic reads, we applied a mixture model to the short-read data, which allowed estimating the relative proportion of each fraction per library. Overall, there was substantial variation in the relative proportions of the three fractions, both among rotifer clones and between libraries of the same clone. Interestingly, the 26% GC-fraction was negatively correlated and the 36% and 48% GC-fractions were positively correlated with genome size based on flow cytometry (FCM) (**Fig. 2b**).

**Fig. 2.**
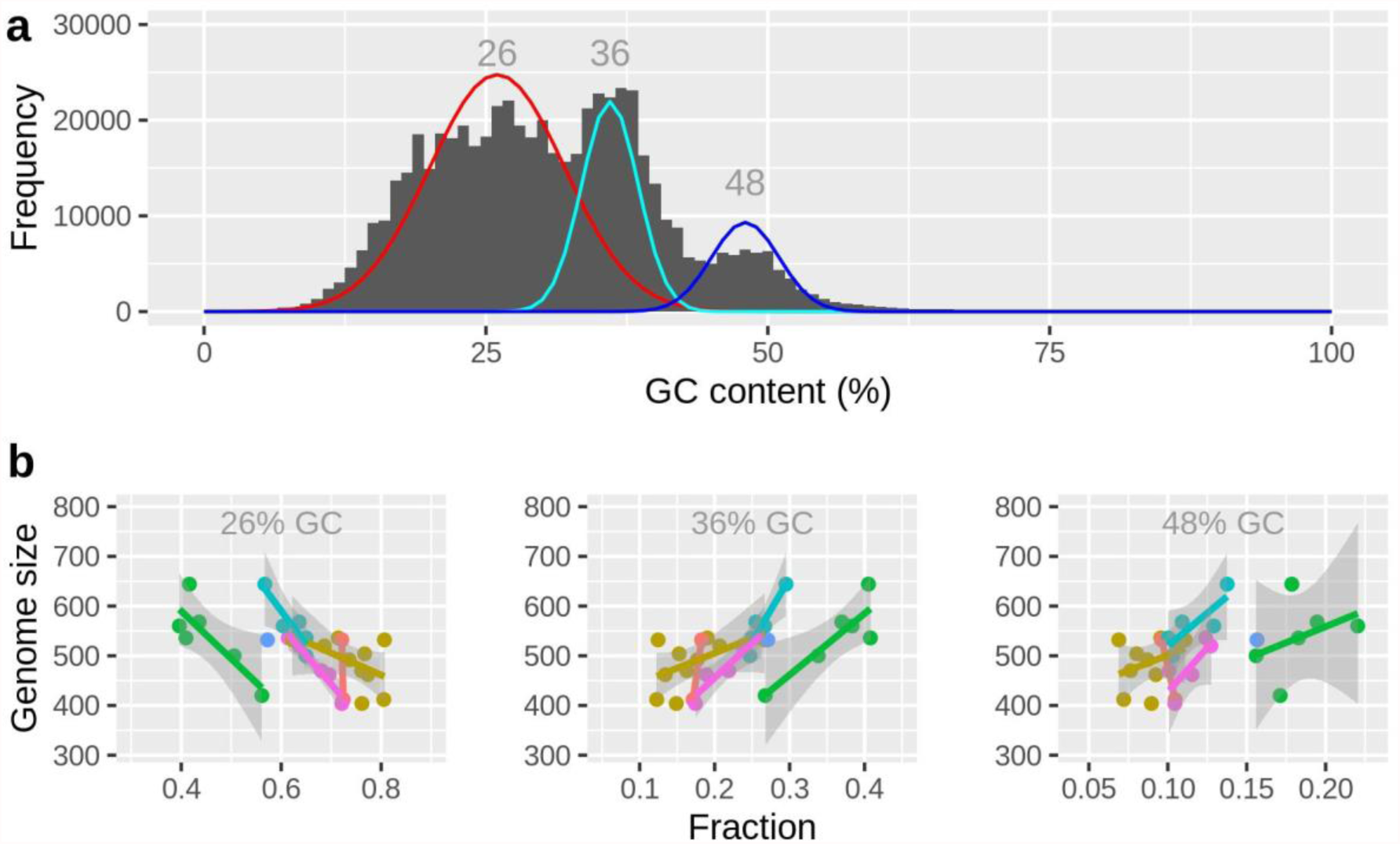
Genome size variation is linked to sequences with elevated GC-content. **a** GC-distribution of all short-read libraries combined (29 libraries of 15 rotifer clones). The red, cyan, and blue lines designate a mixture model fitted to these data, consisting of three normally distributed subpopulations (26 ±6, 36 ±2.5, and 48 ±3 %GC; means and sds). **b** Panels show the results of the same mixture model applied to each library individually, thus estimating the proportion of the total reads per library in each GC-fraction. Genome size estimates are based on flow cytometry and were taken from (Stelzer et al. 2019). Colors in **b** correspond to the six library preps (A-F) listed in Tab. S4: A=orange, B=gold, C=green, D=turquoise, E=blue, F=pink).

We also used two kmer-based tools, GenomeScope 2.0 (Ranallo-Benavidez et al. 2020) and findGSE (Sun et al. 2017), to obtain reference-free estimates of genome size for each clone/library. Those estimates were generally lower than their FCM based counterparts, approximately 0.8-fold in findGSE and 0.6-fold in GenomeScope (**Supplemental file 3**). GenomeScope appeared to struggle at sequencing coverages below ca. 25-fold, where it estimated extremely low genome sizes (compared to FCM) and unrealistically high heterozygosities (6-10%). By contrast, findGSE performed constantly along the gradient of sequencing coverages. Interestingly, there was a positive correlation between genome size (FCM estimate) and the ratio of repeats, a fitted parameter of findGSE (**Supplemental Fig. S4**; Spearman-rank correlation test, ρ=0.609, *P*=0.0057).

Our assembly-based analyses rely on the alignment of cleaned reads of each of the 29 libraries to the reference genome. Total alignment rates (TAR) of reads to the reference genome draft were generally above 94%. TARs were highest in those rotifer clones that are most closely related to the reference genome (**Supplemental Fig. S5**). Concordant alignment rates, i.e., properly aligned reads with the correct insert size, were >90% in all libraries (**Supplemental Table S5**). However, only about 1/3 of the reads aligned uniquely to one site in the genome. Only a small proportion of reads, usually well below 5%, showed discordant alignment, either due to an incorrect insert size, or when only one of the mates aligning to the reference genome. Discordant alignment rates were generally low (1.3-5.1%) and were not correlated to genome size.

To identify CNV regions, we calculated the average per-base coverage for 5kbp and 50kbp windows. To normalize coverage across libraries, we divided the per-base coverage of each window by 1/2 of the (mean) exon coverage of the respective library. This yields a value of 2 for all diploid regions of the genome, corresponding to two copies for those genomic regions (provided that our assembly is unphased in these regions). This analysis of coverage variation revealed that large tracts in the genome of *B. asplanchnoidis* display consistent patterns of coverage variation, which we quantified as the standard deviation of coverage across libraries and clones (**Fig. 3**). For example, the first contig (000F) exhibits large coverage variation with a standard deviation of 1.5-1.7 while the next three contigs (001F to 003F) have much less coverage variation (< 0.5 s.d.) and a mean coverage value of close to two. There are several other contigs showing consistently elevated coverage variation, like 004F, 031F, 032F,036F, 042F, and 044F. In addition, some of these variable contigs appear to be more variable than others (e.g., 031F is more variable than 032F). These overall patterns were very similar when a lower 5kbp-window resolution was applied (**Supplemental Fig. S6**)

**Fig. 3.**
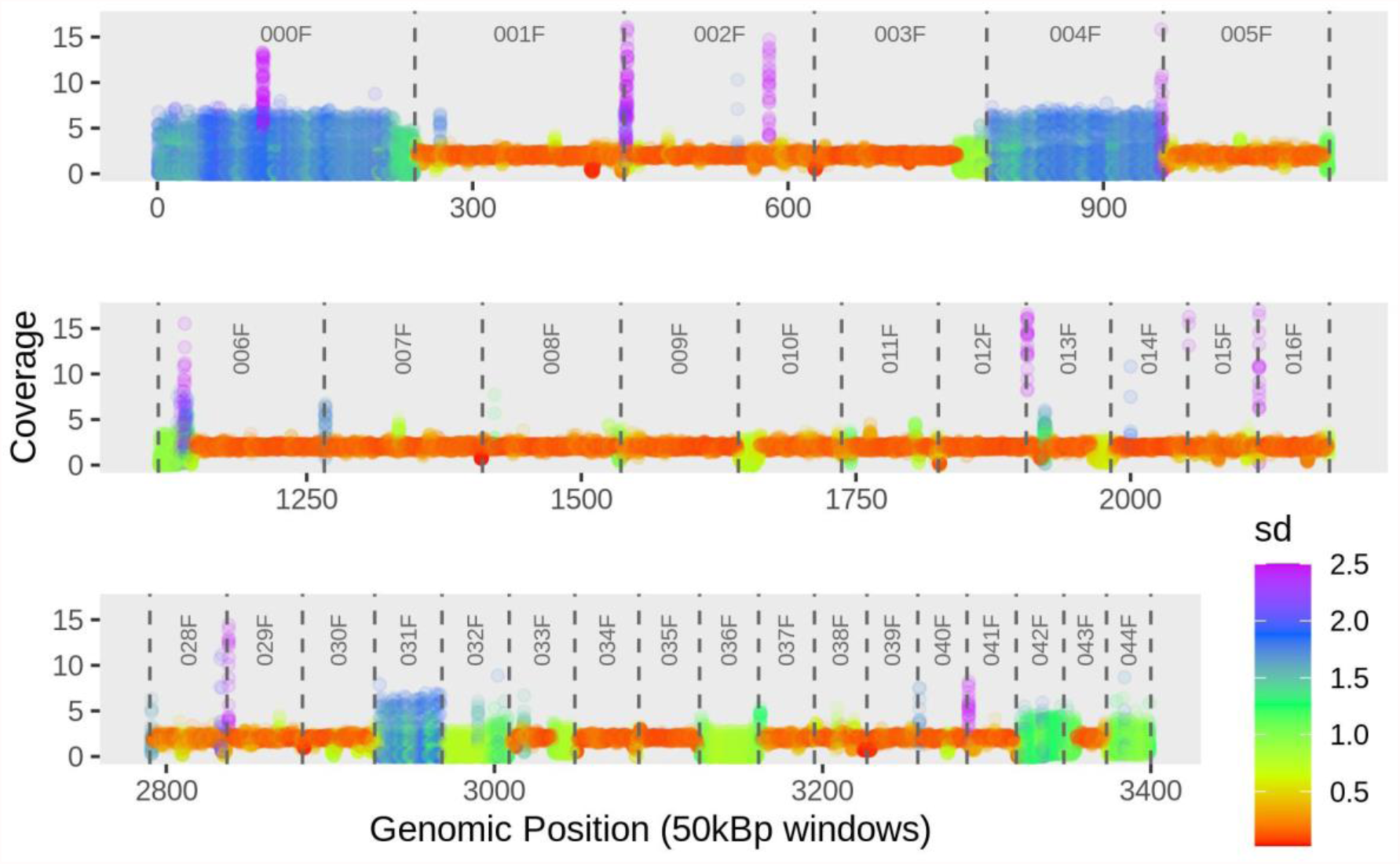
Coverage variation along the *B. asplanchnoidis* genome. This graph shows 34 selected contigs spanning ∼140 Mbp (i.e., about 60% of the assembly). Each circle represents a 50kbp window of one of the 29 sequencing libraries. Coverage (y-axis) was normalized by dividing the per-base coverage of each window by 1/2 of the (mean) exon coverage of the respective library. Contig borders are indicated by vertical dashed lines, and contig IDs are listed on the top. Contigs are ordered by size rather than reflecting biological contiguity.

Variation across clones/libraries is indicated by color (based on standard deviations). Standard deviations larger than 2.5 (about 4% of the data) were capped to a value of 2.5, to allow colored visualization.

Combining the values of coverage variation of all windows (n=4380 for 50kbp, n=45800 for 5kbp) reveals a multimodal distribution with a prominent peak located at low coverage variations of ∼0.15 (lowSD in **Fig. 4a**). This peak corresponds to the genomic sections that are colored in orange/red with a mean coverage of 2 in **Fig. 3**. Interestingly, at intermediate coverage variations (interSD; 0.7 < s.d. < 2.0) there appear to be at least two peaks, which correspond to the green and blue regions in **Fig. 3**, respectively. There are also a few windows showing high coverage variation (highSD; s.d. > 2.0), which form the right tail in **Fig. 4a**.

**Fig. 4.**
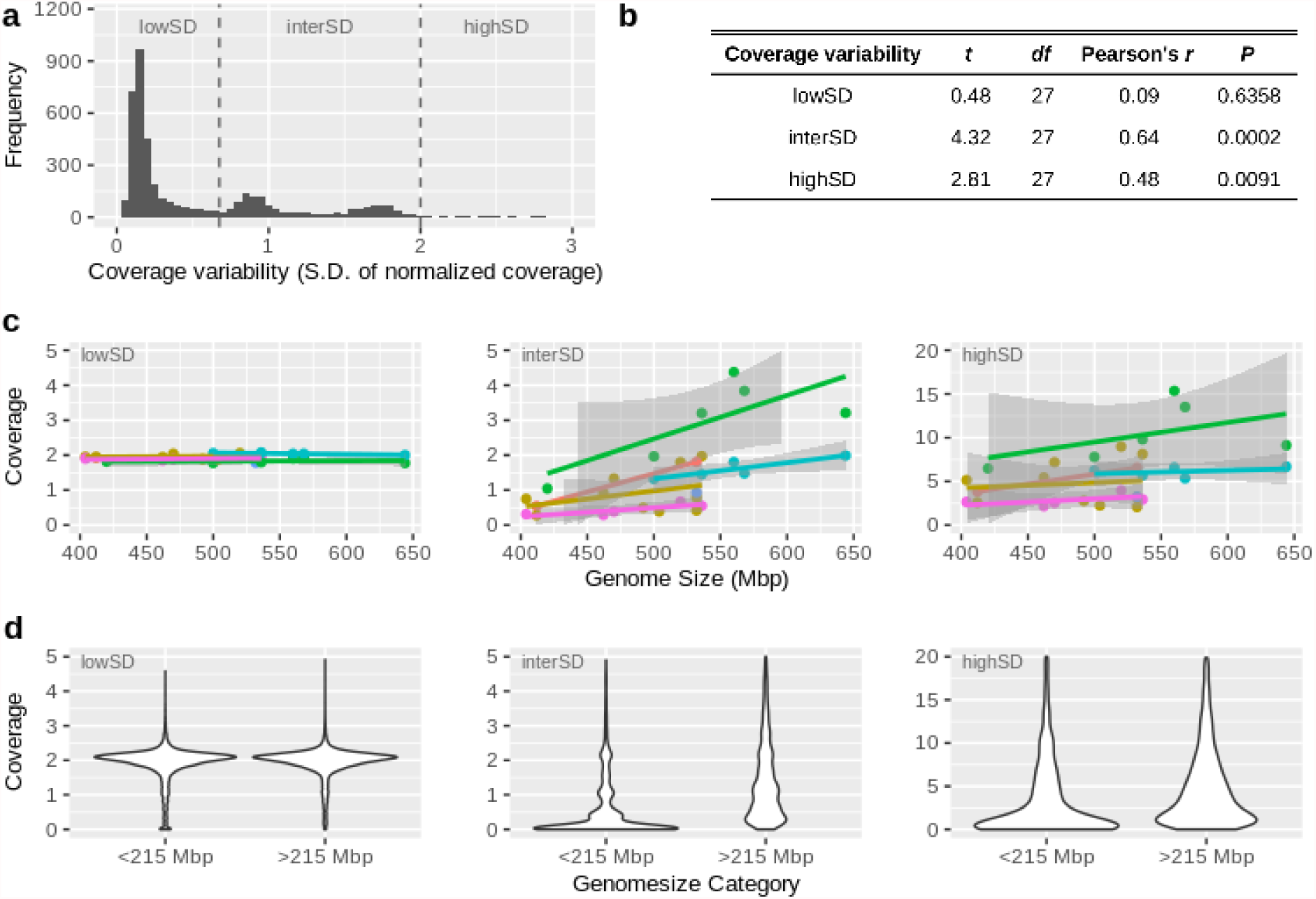
Genomic regions of elevated coverage variability are responsible for genome size variation. The data in this figure is based on normalized coverage (calculated for 50kbp windows) of 29 short-read libraries from 15rotifer clones. **a:** Genomic regions differ in coverage variability (measured as standard deviation of coverage among libraries) as there are regions of low, intermediate, and high variability. **b, c:** Mean coverage in regions of elevated coverage variation (interSD, highSD) correlates positively with genome size (flow cytometry data on genome size was taken from (Stelzer et al. 2019). Dots represent the mean coverage per library. Colors indicate co-prepared libraries (of which some were prepared from the same rotifer clone, but at different dates and with different library preparation methods; see Table S4). Colors in **c** correspond to the six library preps (A-listed in Tab. S4: A=orange, B=gold, C=green, D=turquoise, E=blue, F=pink). **d:** This panel shows the same data as in **c**, but rotifer clones are categorized into small genome sizes (<215Mbp, which represent the basal genome size of *B. asplanchnoidis* according to (Stelzer et al. 2019) vs. large genome sizes (>215Mbp, which contain additional genomic elements that independently segregate during meiosis, according to (Stelzer et al. 2019)).

To test for an effect of these coverage variations on genome size, we calculated the mean coverage for each clone/library for these three categories of coverage variation (lowSD, interSD, highSD) and calculated their correlations with genome size. Notably, there were substantial differences in coverage at the interSD and high SD regions among the different libraries, even in some that had been prepared from the same rotifer clone (**Fig. 4c**). Thus, in order to control for the effect of library preparation, were calculated partial correlations between the variables ‘mean coverage’ and ‘genome size’ (**Fig. 4c**). Those correlations at ‘intermediate’ and ‘high’ variability were highly significant. This result holds even if the libraries with elevated coverage and GC content at interSD and highSD regions (green symbols in Fig. 2b and 4c) are excluded (**Supplemental Fig. S7**)

By merging adjacent 5kbp windows that show increased coverage variation and consistent coverage pattern (significant correlation of coverages), we identified 509 CNV regions in the genome of *B. asplanchnoidis* (**Supplemental file 4**). The total genome space classified hitherto as “copy number variable” was 72.43Mbp (i.e., 31% of the genome assembly). Interestingly, large CNV regions megabases in size made up a large fraction of this total, as can be seen by an “N50” of 0.455 Mbp for the CNV-fraction of the genome. Two of the largest contigs, 000F and 004F, consisted almost entirely of 3-4 large CNV regions, which were separated only by short “breakpoints” of lower coverage variability (**Supplemental file 4**). Our 5kbp window-scanning algorithm also identified CNVs that resided in contigs with otherwise “normal” coverage. In many cases, these CNV regions were near the beginning or end of a contig (e.g., 003F, 006F, 010F). Interestingly, coverage values of clone IK1, a cross between OHJ7i3n2 and OHJ22i3n14, were usually intermediate between those of the two parental clones (Figs. S8, S11, S12). In addition, four clones that were derived by selfing from clone OHJ7 displayed coverages that were largely consistent with their differences in genome size (Fig. S9). For instance, the clone with the largest genome of the selfed line, OHJ7i3n5, had an additional coverage peak at about 2.5 times the base coverage (set by IK1), which indicates that some CNV regions have significantly higher coverage than any other clone of this selfed line.

To additionally classify CNV regions according to their length and contiguity, we considered contigs as “B-contigs” if they contained a large fraction of CNV windows (in analogy to B-chromosomes). Setting this threshold at 90% of contig length, 38 contigs are classified as “B-contigs”, comprising 77% of all CNV-windows, i.e., 55.8 Mbp of the assembly (**Supplemental Fig. S10, S11, Supplemental Table S6**). Thus, approximately three quarters of the observed CNVs affect more or less an entire contig, while the remaining quarter of CNV regions were found on contigs with otherwise low coverage variability.

To independently confirm CNVs, we chose four genomic loci for PCR-amplification, two in CNV regions and two in non-CNV regions (**Supplemental Tables S7, S8**). All four primer pairs yielded amplicons with the correct size, with no signs of non-specific amplification (**Fig. 5a**). The two primer pairs targeted to non-CNV regions (TA_001F and TA_003F) yielded an amplicon in all rotifer clones. In contrast, the two primer pairs targeted to CNV loci (TA_000F and TA_032F) only amplified in some clones. In particular, clones with the smallest genome sizes (OHJ82, OHJ22 and its descendent OHJ22i3n14) apparently lack both CNV loci, and in others (OHJ 98, 104, 105) the TA_000F-locus was present, but the TA_032F-locus was absent. Overall, these patterns were highly consistent with coverage of the amplified regions in sequencing libraries (**Fig. 5c**). Copy numbers for TA_032, as estimated by ddPCR, ranged from zero to six across the studied rotifer clones, including 3 copies in IK1, a cross between OHJ7i3n2 (six copies) and OHJ22i3n14 (zero copies) (**Fig. 5b, Tab. S8**).

**Fig. 5.**
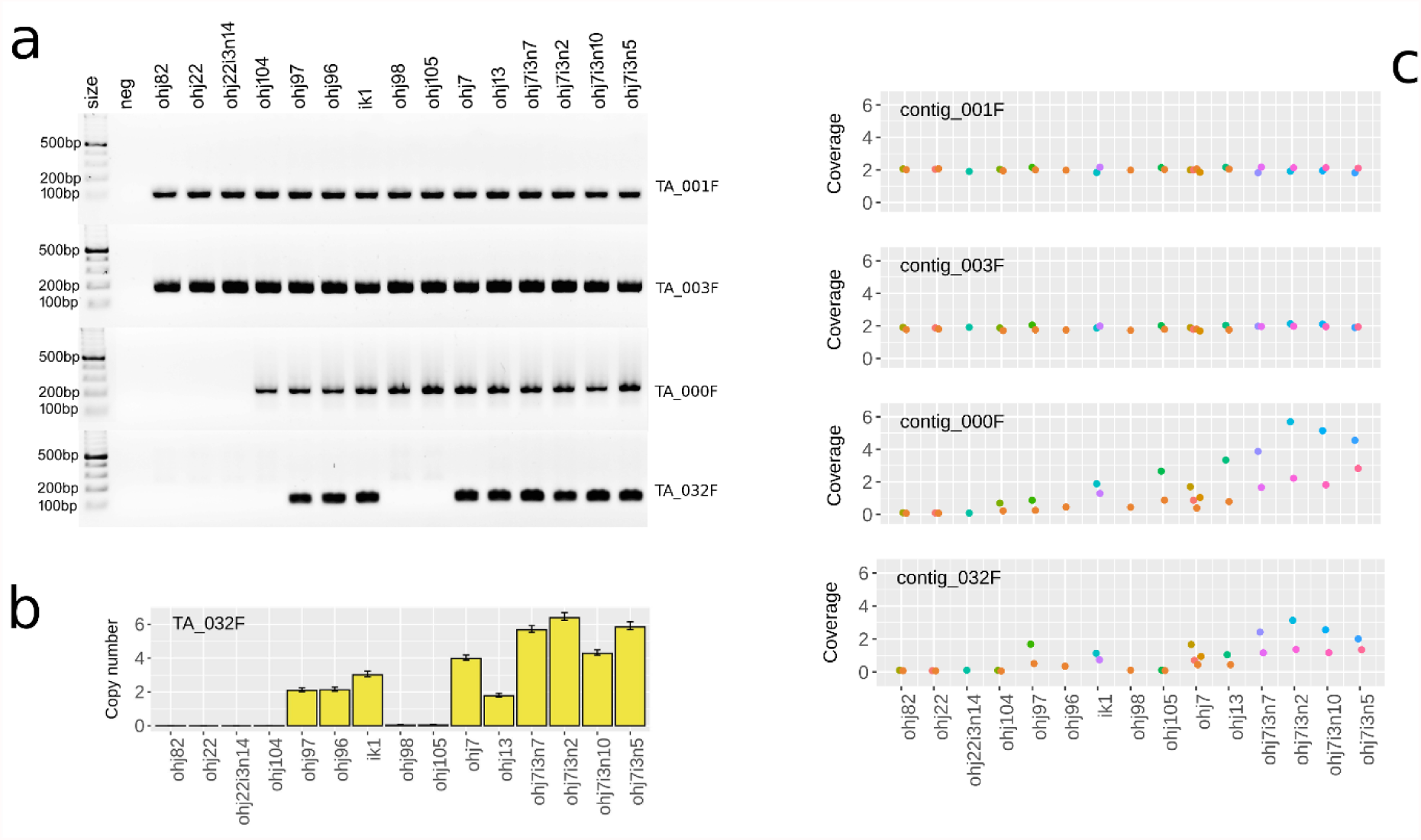
PCR-based confirmation of CNV-loci. All 15 studied clones from the OHJ-population are displayed in order of ascending genome size (from left to right). **a** PCR-products of primers targeting the four candidate loci (non-CNV loci: TA_001F and TA_003F; CNV-loci: TA_000F and TA_032F). **b** Copy numbers of the TA_032F-locus as estimated by droplet digital PCR. Error bars are 95% confidence intervals. **c** Coverage of the amplified region in sequencing libraries of the same 15 rotifer clones (for comparison). Dots are the average coverages across an entire contig. Colors identify co-prepared sequencing libraries.

After having identified the CNV-regions that contribute to intrapopulation genome size variation in the OHJ-population, we annotated repetitive elements of these regions, and compared them to the rest of the genome. A custom repeat library was created using RepeatModeler2, and the top-contributing TEs were curated. In total, 123 Mbp of the assembly (53.6%) were masked by this library. The highest contributing element (rotiSat2) accounts for just over 50 Mbp of this (**Fig. 6a**). The 36 most abundant repeats represent 67% of masked repeats and 82.6 Mbp of the assembly. Of the 36 highly contributing repeats, 7 were enriched in the interSD region, 12 in the highSD region, 5 in allCNVs, 6 in the B90 region, and 5 in the B95 region (**Supplemental file 6**). Overall, repetitive elements, and especially satDNAs were over-represented in CNV-regions (**Fig. 6c, Supplemental file 7**).

**Fig. 6.**
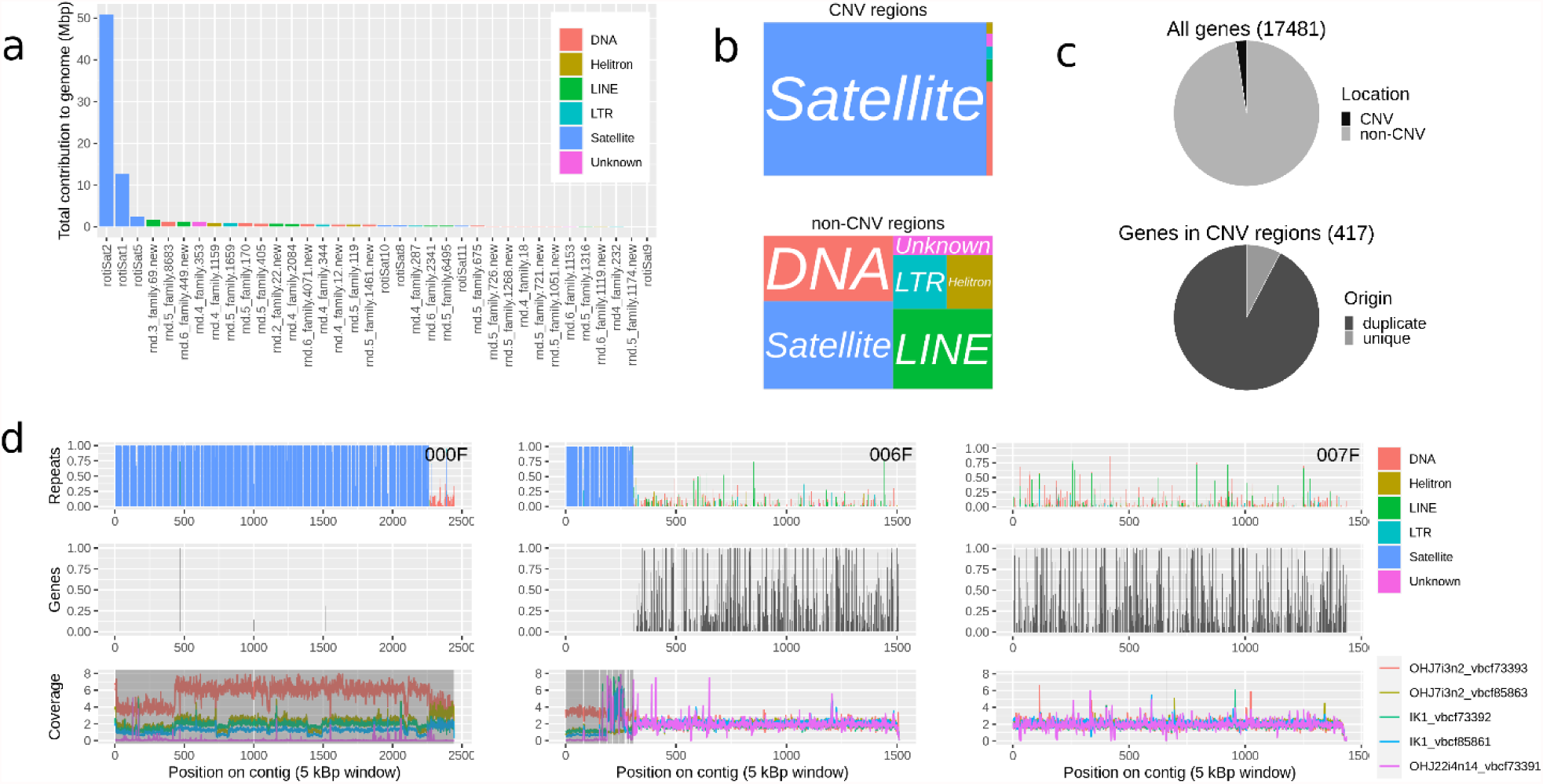
Genome structure of *B. asplanchnoidis*. **a** The top-36 repeat elements ranked according to their contribution to the genome. **b** Differences in repeat element composition between CNV and non CNV regions. **c** Differences in the distribution of gene content between CNV and non CNV regions (top) and the respective proportion of genes derived from gene duplication events (bottom). **d** Three representative examples of contigs: a “B-contig” consisting almost exclusively of CNV regions (000F), a contig containing a large CNV region at one end (006F), and a non-CNV contig (007F). Top panels display the proportion of each 5kb window occupied by repeats, intermediate panels display the proportion of each 5kb window occupied by annotated genes, bottom panels display (exon-)normalized coverages of three rotifer clones (parents: OHJ7i3n2 and OHJ22i3n14, crossed offspring: IK1). Shaded regions indicate CNVs.

Of the satDNAs we identified in B. *asplanchnoidis*, three (rotiSat2, 8 and 9) are not present in any other sequenced *Brachionus* genome; three others (rotiSat1, 5, 10, 11) are shared with the *Brachionus plicatilis* ss. genome. All but two other repeats in the topRE library were found in at least two other *Brachionus* genomes in varying levels. In addition, we identified a DNA/MITE element and an uncharacterized element in the *Brachionus plicatilis* ss. genome that are not found in other *Brachionus* genomes.

CNV-regions differed strongly from non-CNV-regions by having a much lower gene density (**Fig. 6c, Supplemental file 7**). Phylogenetic orthology inference based on proteomes (OrthoFinder analysis) of four species in the *B. plicatilis* species complex, with the bdelloid rotifer *Adineta vaga* as an outgroup, resulted in the assignment of 93063 genes (90.9% of all annotated proteins) to 17965 orthogroups. Fifty percent of genes were in orthogroups with 6 or more genes and were contained in the largest 5228 orthogroups. There were 3953 orthogroups represented in all species and 299 of these consisted entirely of single-copy genes. Many duplication events appear to be species-specific, with an especially high number of genes (5634, or 32 % of all protein coding genes) derived from gene duplication in *B. asplanchnoidis*. While gene density is significantly reduced within CNV regions (417 genes located within CNV regions = 2.39 % of all annotated protein-coding genes, p<0.001, **Fig. 6c**), a significant number of these genes derive from gene duplication events (385 genes = 92.33 % of all genes located within CNVs, *p*<0.001, **Fig. 6c**). The overall pattern of gene distribution thus shows that CNV regions almost exclusively contain gene copies.

GO enrichment analysis of genes throughout the *B. asplanchnoidis* genome that derived from duplication events identified 29 significantly enriched GO terms (**Supplemental Table S9**). When restricting the gene set to only those genes derived from a duplication event that were found within CNV regions, we identified eleven significantly enriched GO terms (**Supplemental Table S10**).

Throughout this study we observed several conspicuous patterns related to GC-content. Regions of elevated coverage variability (i.e., interSD and highSD regions), CNV regions, and B-contigs were characterized by an elevated GC-content showing a main peak at ∼37% GC and two additional peaks at around 50% GC (**Supplemental Figs. S13, S14**). By contrast, regions of low coverage variability had their main peak at ∼25% GC (**Supplemental Fig. S14**). Interestingly, these three peaks were also present in the GC-distributions of unaligned sequencing reads from rotifer clones varying in genome size (Fig. 2b). The 37% GC peak, which was the most prominent peak in genomic regions of elevated SD, in CNVs, and in B-contigs (**Supplemental Fig. S14**) could be attributed mainly to the satellites rotiSat1 and rotiSat2, while the higher peak at ∼55% GC could be attributed to rotiSat5 (**Supplemental Fig. S15-19**). Overall, these three satellites are the most abundant repeat elements in the *B. asplanchnoidis* genome, and their consensus sequences show the same characteristically elevated GC contents compared to most other repeat elements (Supplemental File 6). We also observed one minor but distinct peak at ∼48% GC in highSD regions, CNV regions, and B-contigs, which apparently consisted of sequences that were not classified as repeats by repeatModeler2 (**Supplemental Figures S15-19**).

## Discussion

In this study, we provide a high-quality reference genome draft of the rotifer *Brachionus asplanchnoidis* to shed light on one of the most extreme examples of intraspecific genome size variation in the animal kingdom (Riss et al. 2017; Stelzer et al. 2019). Our genome assembly had a length of 230.1 Mbp. The 2C DNA content of the same rotifer clone is 568 Mbp, according to flow-cytometry based estimates of an earlier study (Stelzer et al. 2019). This earlier study provided evidence that the genome of this particular clone consists of a core haploid genome of 207Mbp and four copies of a segregating 34Mb element. Assuming that our 230.1 Mbp assembly is completely unphased, our reference genome thus amounts to 95% of the flow cytometry-based estimate of the haploid genome (241Mbp).

We aligned short-reads of 15 rotifer clones from the same geographic population, together encompassing a 2C-genome size range of 404–644Mbp, to the reference genome. This analysis uncovered multiple long tracts along the reference genome with increased coverage variation across clones. Additionally, we found that the average coverage at CNV-regions strongly correlates with genome size. CVN-regions also carried a distinct signature in terms of an increased GC-content (36% and 48%, respectively, versus 26% for the rest of the genome), which was both apparent in the short-read data and in the genome assembly. Strikingly, many CNVs had near zero coverage in three of the studied clones (OHJ82, OHJ22, OHJ22i3n14), which was independently confirmed by our PCR-based assays of two selected CNV regions. These results are highly consistent with previous evidence from flow cytometry experiments, showing that these three clones are characterized by a ‘basal’ genome size (i.e., are close do the smallest observed genome size in this species) and that they entirely lack the independently segregating genomic elements (ISEs) that are present in many other members of the OHJ population. Our results indicate the presence of multiple different ISEs in the OHJ population. For example, both our PCR- and alignment data suggested that the two contigs 000F and 032F belong to two different ISEs, because only the former was detectable in the clones OHJ98, OHJ104, and OHJ105 (**Fig. 5**). This observation is consistent with an earlier study, which suggested such diversity based on the size of ISEs measured by flow-cytometry.

CNV regions account for megabase-long tracts in the genome of *B. asplanchnoidis*. Several contigs displayed highly similar coverage patterns across almost their entire length, if one ignores the few and very short breakpoints. Such contigs might be fragments of even larger elements, perhaps B-chromosomes. Contig 032F might be a good candidate for this, ranging from zero to six copies across the OHJ-population, as indicated by ddPCR. We also detected large CNVs, hundreds of kilobases in length, that were located on contigs with otherwise normal (diploid) coverage and low coverage variation (e.g. contig 006F in **Fig. 6d**). Such a genomic pattern is consistent with a stable diploid chromosome that contains a large homozygous insertion in some clones, hemizygous insertions in others, and a homozygous deletion in the remaining clones. Additionally, there might be length polymorphisms in such genomic regions, which are dominated by tandemly repeated satellite DNA. In the future, assembling multiple genomes from rotifer clones with different genome size, ideally using long-read technologies in order to approach chromosome-level assemblies (Simion et al. 2020), might allow a more precise delineation of individual CNVs into these two categories. Overall, our data is consistent with a mixture of B-chromosomes and large insertions into normal chromosomes, and possibly a dynamic exchange between both genomic fractions (since they are made up mostly by the same set of tandemly repeated satellite DNA; see below). Interestingly, GO enrichment analysis of genes that derived from a duplication event identified one term, GO:0015074 (DNA integration), as the most significant term (*P* < 1e-30), which might indicate elevated transposon activity during the early evolution of the *B. asplanchnoidis* genome.

There are a few technical caveats and limitations to be considered in our analysis. First, although we found strong correlations between coverage variation at CNV regions and genome size variation, it was not possible to quantitatively “predict” the genome size of individual clones based on coverage along the reference assembly. Library preparation method seemed to introduce additional variation, specifically at the CNV regions, that prevented us from determining exact copy numbers of these genomic regions. This is a well-known limitation of short-read libraries from genomes that contain large amounts of satellite DNA (Lower et al. 2018), in particular when combined with heterogeneous GC contents (Benjamini and Speed 2012). We could alleviate this limitation and determine exact copy numbers by using digital droplet PCR, finding that the targeted genomic region (contig 032F) is present in zero to six copies across the OHJ-population. This approach is a promising strategy for future studies to accurately assess copy number variation across many loci and in a large number of genomes from the same population. In addition, the differences in copy numbers at CNV regions could be estimated by comparing clones within a set of library preparations. For instance, if coverage at CNV regions is scaled by a reference clone (like clone ‘IK1’ in Fig. S8 and S9), coverage at CNV regions of focal clones tends to fall into discrete clusters, indicating n-fold differences in coverage relative to the reference. A second limitation is that our reference genome might still not contain all the ISEs that contribute to genome size variation in the natural OHJ population, simply because it represents just a sample from this population. To obtain a more complete picture, more genomes of the OHJ-population would be needed to be sequenced, preferably using long-read sequencing technologies that allow better identification of structural variants (De Coster et al. 2019). Third, one might argue that our CNV-detection pipeline could miss many of the smaller insertions and deletions, in particular those smaller than 5 kilobases. However, it is quite unlikely that such small-scale structural variation has much influence on genome size variation in *B. asplanchnoidis*, since the percentage of discordantly aligned reads was overall rather low (1.3-5.1%) and it did not scale with genome size. With our reference genome being at the upper end of the genome size distribution of the OHJ-population, we would expect to find a higher percentage of discordant reads in smaller genomes (mainly due to deletions), which was not the case.

CNV regions differed strongly from the remaining genome regarding repeat composition and gene density. Strikingly, most CNV regions were composed of only three satellite repeat elements, some unique to *B. asplanchnoidis* and others shared only with its closest congener, *B. plicatilis*, indicating recent evolutionary origin. The two most prominent satellite repeats, rotiSat1 and rotiSat2, consisted of 154 and 143bp monomers that were tandemly repeated for up to a few megabases. The low sequence diversity of CNV regions nicely explains the characteristic trimodal GC-content signature mentioned earlier, especially since the most abundant satellite elements display the same elevated GC content (**Supplemental file 6**). Non-CNV parts of the genome contained a much higher diversity of other repeat elements, which included DNA transposons, LINEs, LTR elements, and Helitrons. Gene density was significantly lower in CNV regions compared to the rest of the genome, and genic regions in CNVs were typically confined to short stretches scattered across the contig. Interestingly, Orthofinder suggested that genes in CNV regions were three times more likely to derive from a duplication event than genes found in other places of the genome. This indicates that duplications had a role in the origin of these CNV-regions, which incidentally has some resemblance to the proposed early evolution of B-chromosomes (Ahmad and Martins 2019; Ruiz-Ruano et al. 2019).

In this study, we have identified, for the first time, genomic elements that can cause substantial within-population genome size variation, ultimately allowing investigation of genome size evolution at microevolutionary time scales. In the OHJ-population of *B. asplanchnoidis*, these genomic elements consist of up to megabases-long arrays of satellite DNA, with only few interspersed genes or other sequences. Though satellite arrays can form essential chromosome structures such as centromeres and telomeres (Garrido-Ramos 2017), the large intrapopulation variation of these elements in *B. asplanchnoidis*, and their virtual absence in some individuals suggests that, overall, these DNA additions do not provide an immediate fitness benefit to their carriers. Nevertheless, variable amounts of ‘bulk DNA’ might influence the phenotype through subtler mechanisms independently of the DNA information content, for example through potential causal relationships between genome size and nucleus size, or cell size (Gregory 2001; Cavalier-Smith 2005). Increased levels of structural variation in a genome, as implied by our findings, may also constrain adaptive evolution or genome stability over microevolutionary time scales. In this regard, the genome of *B. asplanchnoidis* should be a valuable addition to existing models of genome evolution, enabling whole-genome analysis combined with experimental evolution approaches (Fussmann 2011; Declerck and Papakostas 2017).

## Methods

### Origin of rotifer clones and DNA extraction

Resting eggs of rotifers were collected in the field from Obere Halbjochlacke (OHJ), a small alkaline playa lake in Eastern Austria (N 47°47′11″, E 16°50′31″). Animals were kept in clonal cultures. A rotifer clone consists of the asexual descendants of an individual female that hatched from a single resting egg. Since resting eggs are produced sexually in monogonont rotifers, each clone has a unique genotype. Our clones from the OHJ-population have been characterized previously with regard of their genome size and other biological traits (Stelzer et al. 2019).

Rotifers were cultured in F/2 medium (Guillard 1975) at 16 ppt salinity and with *Tetraselmis suecica* algae as food source (500–1000 cells μl^−1^). Continuous illumination was provided with daylight LED lamps (SunStrip, Econlux) at 30–40 µmol quanta m^−2^ s^−1^ for rotifers, and 200 µmol quanta m^−2^ s^−1^ for algae. Stock cultures were kept either at 18 °C, re-inoculated once per week by transferring 20 asexual females to 20ml fresh culture medium, or they were kept for long-term storage at 9°C, replacing approximately 80% of the medium with fresh food suspension every 4 weeks.

To produce biomass for DNA extraction, rotifers were cultured at 23°C in aerated borosilicate glass containers of variable size (250mL to 20L). Prior to DNA extraction, rotifers were starved overnight in sterile-filtered F/2 medium, with 2-3 additional washes with sterile medium on the next day. In most preparations we also added the antibiotics Streptomycin (Sigma-Aldrich: S6501) and Ampicillin (Sigma-Aldrich: A9518) to the washing medium, both with an end concentration of 50mg/ml. DNA was extracted using the Qiagen kits Dneasy (for short-read sequencing; from approximately 5000-7000 rotifers) and GenomicTips 100 (for long read Pacbio sequencing; from >20000 rotifers), and RNA was extracted from freshly prepared biomass with Rneasy.

### Sequencing of the reference clone

We selected one rotifer clone (called: OHJ7i3n10) as DNA-donor for the reference genome. This clone ultimately derives from an ancestor of the natural OHJ population. However, its immediate ancestors were passed through three generations of selfing (i.e., mating one male and female of the same clone). More details on the genealogy of this lineage and its biological characteristics can be found in (Stelzer et al. 2019). According to this study, OHJ7i3n10 has a 2C-genome size of 568 Mbp, and thus contains approximately 40% excess genomic sequences, compared to the smallest genome size of the OHJ-population (∼410 Mbp).

Our *B. asplanchnoidis* reference genome is based on long-read sequencing technology (PacBio SMRT® on the Sequel-platform). In total, we obtained 16.3 Gbp from two SMRTcells, which confers to 57-fold coverage assuming a haploid genome size of 284 Mbp. Additionally, we obtained 35.5 Gbp of Iso-Seq transcriptome data and 12 Gbp of short-read Illumina data of OHJ7i3n10. All sequencing and library preps related to the reference genome were performed by the Next Generation Sequencing Facility at Vienna BioCenter Core Facilities (VBCF), member of the Vienna BioCenter (VBC), Austria.

### Sequencing of individuals of the OHJ-population

To characterize genomic variation across the OHJ-population, we generated short-read sequencing data (Illumina platforms HiSeq and NextSeq) from 15 different rotifer clones, both from the natural OHJ population and from various clones of a selfed lineage (**Supplemental Table S4**). Clones were selected such that they covered the full range of genome size of the OHJ-population. Short-read libraries were constructed either using the KAPA HyperPrep kit (Roche), or the NGS DNA Library Prep Kit (Westburg). The KAPA library preparations were done at the Marine Biological Laboratory (for more details, seeBlommaert et al. 2019) while the Westburg preparations were done at VBCF. The two methods mainly differ in the fragmentation method (ultrasonic fragmentation in KAPA *vs*. enzymatic fragmentation in Westburg). Both methods are claimed to deliver sequence-independent fragmentation and to yield consistent coverage across a wide range of GC-contents. While the peak fragment size was ∼400bp in both methods, we observed that the libraries constructed with the Westburg kit sometimes had a pronounced right tail with some fragments up to ∼2000bp. We accounted for these larger fragments by adjusting the relevant parameters during short-read alignment (see below). In many of our clones, we used both library construction methods, yielding in a total of 29 libraries (**Supplemental Table S4**).

### Reference genome assembly and annotation

Pacbio sequences of the reference genome were assembled using the HGAP4 pipeline (Chin et al. 2013) at VBCF, and contamination was initially checked using CLARK (Ounit et al. 2015) against all available bacterial genomes from NCBI (**Supplemental file 1**). We polished this initial VBCF assembly with short-read Illumina data of the identical *B. asplanchnoidis* clone using Pilon (Walker et al. 2014) in three rounds. To investigate the assembly quality, we backmapped the Illumina data to the genome assembly using bwa mem (Li and Durbin 2009) and calculated summary statistics using QUAST (Mikheenko et al. 2018) and Qualimap (Okonechnikov et al. 2016). To provide an additional check for contamination, we used Blobtools (Laetsch and Blaxter 2017) based on the backmapping alignments and a blastn search against the nt database (NCBI).To assess the assembly’s completeness, we performed a BUSCO v4.0.6 (Simão et al. 2015) using the metazoan gene set (n=978) in the *genome* mode applying the *–long* option.

Genes were structurally annotated on the repeat masked genome assembly using the MAKER2 annotation pipeline (v2.31.10, Cantarel et al. 2008) using evidence from Pacbio Iso-Seq transcripts of the same *B. asplanchnoidis* clone; protein homology evidence from the UniProt database (download January 2020, The UniProt Consortium 2017) in combination with proteoms of different clones and closely related *Brachionus* species (**Supplemental Tab. S1**;, and *ab initio* gene predictions from SNAP (Korf 2004), GeneMark-ES (v4.48_3.60_lic, Brůna et al. 2020) and Augustus (v3.3.3, Stanke et al. 2006). The SNAP model was initially trained on the genome assembly with additional support of BUSCO complete hits (see above). GeneMark was computed in the ES suite on the soft masked genome assembly. The ab initio training of the Augustus model was computed on the genome assembly supported by the unassembled Iso-Seq transcripts. In order to compensate for underrepresented rotifer protein representation in the UniProt database, we combined this dataset with proteomes of another *B. asplanchnoidis* clone and four closely related *Brachionus* species (**Tab. S1**). Completeness of these proteomes was checked with BUSCO in the *protein* mode and missing orthologs were compared to determine if completeness could be increased through the combination of proteomes. The combined proteome finally contained 97 % of the BUSCO genes of the metazoan dataset.

MAKER was run over three rounds. For the first round of MAKER, we only used the Iso-Seq data as EST evidence and the combined UniProt and proteome sequences as protein homology evidence applying default parameters and est2genome=1, protein2genome=1 to infer gene predictions directly from the transcripts and protein sequences. We used the gene prediction of round 1 to retrain the Augustus and SNAP model, since the GeneMark model was exclusively computed on the genome assembly and did not require retraining. Maker was run in the second round using the retrained models and again est2genome=1, protein2genome=1 in order to increase prediction fidelity. After a second round of retraining the Augustus and SNAP model, MAKER was run through round 3 with switched off est2genome and protein2genome inference. After structural annotation via MAKER, we functionally annotated the genes using functional classification of genes from InterProScan (Jones et al. 2014) in combination with putative gene names derived from Swissprot (Boutet et al. 2016).

To examine orthologous genes among rotifer species and identify gene duplication events, we used Orthofinder (Emms and Kelly 2019). This analysis was based on proteome information of the *Brachionus plicatilis* species complex: *B. rotundiformis*, *B.* sp. ‘Tiscar’ and *B. plicatilis* (respectively “Italy2”, “TiscarSM28”, “Tokyo1” in, Blommaert et al. 2019), and *B. asplanchniodis* (annotation of this study). Proteome information of *Adineta vaga* (Flot et al. 2013) was included as outgroup. We extracted the longest transcript variant per gene to avoid duplicates in the input proteomes and followed the manual instructions of Orthofinder. To analyze the genomic distribution of genes which were identified to derive from gene duplication events, we extracted all genes of the *B. asplanchniodis* node (Orthofinder: SpeciesTree_Gene_Duplications), removed duplicates to create a non-redundant list of genes. Comparing this list of genes to genes that are located inside or outside of genomic regions with high levels of CNVs, allowed the estimation of non-random gene distribution patterns via Monte-Carlo permutation tests (1000 permutations).

To characterize the putative functional properties of genes derived from a duplication event, we performed gene ontology (GO) enrichment analyses on the set of all multiple copy genes (n=5634) and on the more exclusive set of duplicated genes found within CNV regions (n=385). The reference list consisted of 10083 protein-coding genes (60 % of all annotated genes) with GO annotation via INTERPROSCAN (see above). Enrichment analysis was done with the topGO R package (v2.24.0, Alexa and Rahnenfuhrer 2016) in the category ‘biological process’ using the weight01 algorithm and Fisher statistics (significance level p<0.05).

### Annotation of repetitive elements

The repeat library was produced using RepeatModeler2 (default settings, Flynn et al. 2020) and the polished PacBio assembly. This produced 484 consensus sequences, 289 of which are “Unknown”. These consensus sequences were then used to mask the genome assembly using RepeatMasker with default settings. From this, we identified top contributing repeat elements and used their consensus sequences were used as blast queries against the genome assembly, and the top hits for each consensus were used to produce alignments for manual curation (Platt et al. 2016) and classification (Wicker et al. 2007). Satellite monomers were identified using Tandem Repeat Finder (Benson 1999). Consensus sequences of the satDNAs are dimers of these identified monomers.

Contributions per repeat were estimated by summing up the total length covered by each copy in the RepeatMasker output file. Top contributions were calculated over the whole assembly, and over regions of the genome with distinct coverage variability (lowSD, interSD, and highSD in Fig. 4). For each region, the top 20 repeats were included, resulting in a total of 38 repeats to curate.

Curation was done by manually inspecting each alignment, identifying the ends of the aligning regions, and producing a new consensus sequence. Additionally, classification was performed by searching for TSDs, TIRs, LTRs, satellite structure, and conserved domains. RepBase searches were rarely constructive since rotifer, and especially monogonont, TEs are not well-represented in any databases. The final curated library of topREs contains 37 consensus sequences from 36 elements (one sequence was removed because alignments were to only scattered AT-rich regions, one was an rRNA gene, and one LTR sequence was split into the LTR and internal portions). Redundant sequences between the topRE library and the RepeatModeler library were identified using RepeatMasker. Sequences which were at least 95% identical and covered at least 80% of the uncurated repeat were removed from the uncurated repeats prior to merging the libraries. Due to the short length and difficulties in automatically creating consensus sequences for satDNA elements, all uncurated elements that were identified as similar by RepeatMasker, regardless of similarity or length of hit, were aligned to the curated consensus sequence and visually inspected for alignment. Uncurated elements which aligned across most of the curated satDNA consensus sequences were removed. The combined library (Supplemental file 8) of curated and uncurated elements contained 472 elements (262 unknowns), and was used to mask the *B. asplanchnoidis* genome and to repeatmask related, high quality *Brachionus* genome assemblies to identify shared elements (Kim et al. 2018; Han et al. 2019; Park et al. 2020). For each of the top contributing elements and each genomic region, an enrichment index was calculated as the proportion of each repeat contribution in that region vs the whole genome (i.e. repeat contribution in region/total contribution of repeat) divided by the proportion of the genome represented by each region (i.e. region size/ assembly size). If this index was over 1, it meant that the repeat in question was enriched in the region in question.

### Preprocessing of short-read data

Trimming and adaptor removal was done using bbduk (v38.34, https://jgi.doe.gov/data-and-tools/bbtools/) with the settings: k=23 ktrim=n mink=11 hdist=1 tpe tbo qtrim=rl trimq=20 maq=10 minlen=40. Trimmed reads were initially aligned to the reference genome using bowtie2 (Langmead and Salzberg 2012). Preliminary tests with various parameter settings (local *vs*. end-to end alignment, different settings for the sensitivity parameter, map as single reads *vs*. map as paired reads) did not indicate strong differences. Thus, we used the default values for most parameters, except for fragment size (parameter: X), which was set to the maximum observed fragment size of each library instead of the default value of 500 bp (see **Supplemental Tab. S4**).

We included three steps to remove contaminants, since DNA extracted from the whole bodies of microscopic organisms might contain DNA from other organisms, such as bacteria. First, we extracted all unmapped reads from an initial alignment to the reference genome and screened only those for potential contaminant reads. Thus, we considered the mapped fraction as ‘rotifer DNA’, since they mapped to the (contaminant-free) assembly. Second, the unmapped reads of all libraries were combined and assembled using meta-genomic approaches. For this assembly, we used metaVelvet (v1.2.02, Namiki et al. 2012) with a kmer-legth of 101bp, an insert size of 500 (+/- 200 standard deviation), and a minimum reported contig length of 300bp. The resulting assembly was then analyzed with metaQuast (Mikheenko et al. 2016) using the option ‘automatic pulling of reference sequences’, which restricts the search to bacteria and archaea. Subsequently, the complete genomes of putative contaminants were downloaded, and the unmapped reads were mapped against those genomes. In the third step, the remaining fraction (i.e., reads not mapping to the microbial metagenome) were subjected to a kmer-based identification using the kraken2 pipeline (Wood et al. 2019), with the databases bacteria, archaea, viral, UniVec-Core, and protozoa. We also performed checks on the false-discovery rate of kraken2, by running the same pipeline on reads that initially did map to the reference genome. Finally, all unmapped reads that could *not* be taxonomically assigned to contaminants, with either of the two approaches above, were considered to be of ‘rotifer origin’ and were merged with the mapped fraction. Those ‘rotifer reads’ were then further cleaned by removing mitochondrial DNA, which was done by mapping them to the published mitochondrial genome of *B. plicatilis* (Suga et al. 2008), and by removal of duplicates using FastUniq (Xu et al. 2012). These ‘final reads’ were again mapped to the reference genome, or they were analyzed using reference-free approaches that do not require alignment.

### Analysis of unaligned reads

‘Final reads’ were subjected to kmer-based analyses using a kmer-size of 21bp with jellyfish (v2.1.4, Marçais and Kingsford 2011). Then GenomeScope 2.0 (Ranallo-Benavidez et al. 2020) and findGSE (Sun et al. 2017) were used to obtain kmer-based estimates of coverage, heterozygosity and genome size. Per-base coverages *C* were computed from the kmer-coverages

*C*_k_ with the formula:

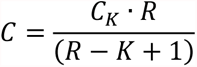

where *R* is the average read size, obtained from dividing the total number of basepairs in each library by the total number of reads. The coverage estimates from these two programs were contrasted with the naïve coverage estimate, based on sequencing effort (total number of bp in a library) and 1C genome size estimated from flow cytometry, assuming a diploid genome.

To analyze GC-distributions among short-read libraries, the GC contents of individual reads were extracted using the function fx2tab of SeqKit (Shen et al. 2016). The resulting csv-files were further analyzed with the R-package mixtools (Benaglia et al. 2009) using the function normalmixEM.

### Analysis of copy number variation

Analysis of copy number variation was done separately for all 29 short-read libraries from 15 different clones of the rotifer *B. asplanchnoidis*. Average per-base depth-of-coverage (DOC) values along 50-kbp and 5kbp windows, respectively, were extracted from the BAM alignment files (‘final reads’ to reference genome) using the samtools function ‘bedcov’ (Li et al. 2009). In total, there are 4835 windows at 50kbp resolution, and 46255 windows at 5kbp resolution in the current genome assembly. To allow comparisons among clones and libraries, DOC was normalized by dividing the per-bp coverage of each window by 1/2 of the (mean) exon coverage of the respective library. In unphased sections of the genome assembly, we expected DOC values of around 2, provided that both alleles of a (diploid) genome map to the correct location. Coverage variation was quantified as the standard deviation of DOC per window (50kbp or 5kbp) across all 29 libraries.

To identify individual CNVs and to locate possible breakpoints within contigs, we used a custom-written R-algorithm involving the following criteria for merging adjacent windows (5kbp) based on the similarity of coverage patterns. First, the coverage variation (measured as the standard deviation of per-base normalized coverage across all libraries) had to be above a defined threshold (i.e., 0.7, which was an *a-posteriori* determined threshold). Second, the read-depths of each library of both windows had to be significantly correlated with each other at the *P*<0.05 level. This was done by calculating the partial correlation coefficients. If both conditions applied, those two windows were considered to belong to the same CNV. The very first and the last window of each contig, which often showed deviant coverage patterns, were merged with their neighboring window. Third, adjacent CNVs identified according to the above criteria were merged if they were separated by only one window (CNV stopping breakpoint) AND if the coverages in the two windows surrounding the breakpoint are significantly correlated with each other. Finally, we only considered CNVs with lengths of at least three adjacent windows. Thus, we obtained a table of all CNVs along the genome of *B. asplanchnoidis*, together with their size, and their location on individual contigs (i.e. in the middle of the contig, at the edges, or spanning the entire contig). Data analysis related to coverage variation and CNV detection was done using custom-written algorithms in the R environment (R Development Core Team 2020) with the base package (v3.6.3) and the add-on packages *stringr* (Hadley 2019) and *reshape2* (Hadley 2007). For graphical visualization, we used ggplot2 (Hadley 2016) and the add-on packages cowplot (Wilke 2020) and treemapify (Wilkins 2021).

### PCR-confirmation of CNV regions

To independently confirm presence or absence of CNV regions in different rotifer clones, and to estimate the copy numbers of these genomic regions, we used PCR-based methods. To identify unique PCR-primer binding sites in in regions of high and of low coverage, we used Thermoalign (Francis et al. 2017) searches in multiple regions of 5000 bp length, spread across the genome and on different contigs. The exact search parameters for Thermoalign are given in **Supplemental file 5**.

Candidate primers were tested and optimized using PCR on template DNA from different OHJ clones, including the reference clone OHJ7i3n10. PCR reactions consisted of 25µl HotStart Taq master mix (Qiagen), 0.1µM Primer, 3mM MgCl2 and 20 ng template DNA. PCR cycling conditions were: 95°C for 15 min, 30 cycles of 94°C for 20 sec, 56°C for 20 sec, and 68°C for 10sec; followed by 68°C for 5 min and hold at 4°C. Agarose gels were used to test for presence/absence of the associated loci across different members of the OHJ population. In total we screened four loci located on different contigs of the reference genome assembly. Two of them were located in copy number-invariable, diploid regions of the genome (according to the coverage estimates from the short-read alignments) and two were located in CNV-regions (**Supplemental Table S7**). In addition to the qualitative PCR test, we used droplet-digital PCR (ddPCR) to estimate the copy number of the CNV loci for each rotifer clone. For each 22µl reaction we used EvaGreen Supermix (Biorad), 0.15µM Primer, 4 units EcoRI and template DNA equivalent of 1500 genome copies. PCR cycling conditions were: 95°C for 5 min, 40 cycles of 95°C for 30 sec, and 61°C for 1 min with a ramp rate of 2°C/sec; 4°C for 5 min, followed by 90°C for 5 min and hold at 4°C. For droplet generation and fluorescence readout, we used a QX200 Droplet Generator and Droplet Reader (Biorad), respectively. Copy numbers (*CN*) of CNV loci were estimated for each rotifer clone using the ratios of amplicon molecule concentrations, which were obtained with the QuantaSoft Software (Biorad):

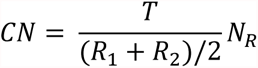

where *T* is the amplicon concentration of the target locus (the one showing copy number variation across the OHJ population), *R*_1_ and *R*_2_ are the amplicon concentrations of the two reference loci, and *N_R_* is the number of copies of the reference loci in the genome (in this case, *N_R_* = 2, since the reference loci were both diploid). The 95% confidence intervals obtained from QuantaSoft were used as an indication of the measurement error.

## Acknowlegements

We thank Hermann Schwärzler for bioinformatic support in establishing the Thermoalign pipeline. Pia Scheu and Martina Nindl provided valuable advice for optimization of ddPCR conditions. The participants of the “Uppsala TE Jamboree” provided support in classifying repetitive elements.

Funding for this project was provided by the Austrian Science Fund, grant number P26256 (PI: CPS), and by a scholarship for the promotion of young researchers by the University of Innsbruck to JB. JB is currently supported by funding from SciLifeLab, Uppsala.

## Supplementary Information

### This file includes

Supplementary notes on short-read preprocessing

Figures S1 to S19

Tables S1 to S8

References for SI citations

### Data to be submitted as electronic supplementary files

Supplementary file 1 (VBCF report on genome assembly and contaminant filtering; pdf-file)

Supplementary file 2 (Summary of short-read preprocessing and fastqc-reports; xlsx-file)

Supplementary file 3 (Kmer-based analysis of cleaned Illumina reads, xlsx-file)

Supplementary file 4 (Ranges of all CNVs across the *B. asplanchnoidis* genome; csv-file)

Supplementary file 5 (Input parameters of the thermoalign pipeline; txt-file)

Supplementary file 6 (Detailed information on top-36 contributing repeat elements; xlsx-file)

Supplementary file 7 (Repeat profile, Gene density, and CNVs of the 50 largest contigs; html-file)

Supplementary file 8 (Combined library of curated and uncurated repeat elements; fasta-file)

### Data to be submitted to public databases

- Raw reads
- Assembly
- Gene annotation files
- Repeat annotation

### Supplementary notes on short-read preprocessing

In total, we obtained 265.6 Gbp of Illumina short-read data from the 15 different *B. asplanchnoidis* clones. After all preprocessing steps had been completed, this amount was reduced to 194.3 Gbp. Total alignment rates to the reference genome improved from 90.9-98.4%, after quality-filtering only, to 94.1-98.4%, after contaminant removal (**Supplementary file 2**) Contamination rates were variable among the different libraries (**Fig. S1**). Contaminant DNA mostly derived from bacteria, of which *Pseudomonas toyotensis* was the only bacterial genome that could be identified with our metagenomic assembly-based approach. Among the libraries 3-30% of the unmapped reads could be assigned to other microbial contaminants. Contamination by the food algae *Tetraselmis* was low to absent depending on library (usually less than 1% of unmapped reads), and protozoan contamination was below the detection threshold in all libraries. After preprocessing, the genome coverages ranged between 9.4-79.0-fold, if the Gbp sequence of each library is expressed as multiples of the 230.1Mb *B. asplanchnoidis* reference genome, with the majority of libraries being above 20-fold (**Supplementary file 2**). Quality control using fastqc [1] indicated substantial improvements in most parameters in the course of the pre-processing pipeline, with the notable exception of GC-content (**Supplementary file 2**)

**Fig. S1:**
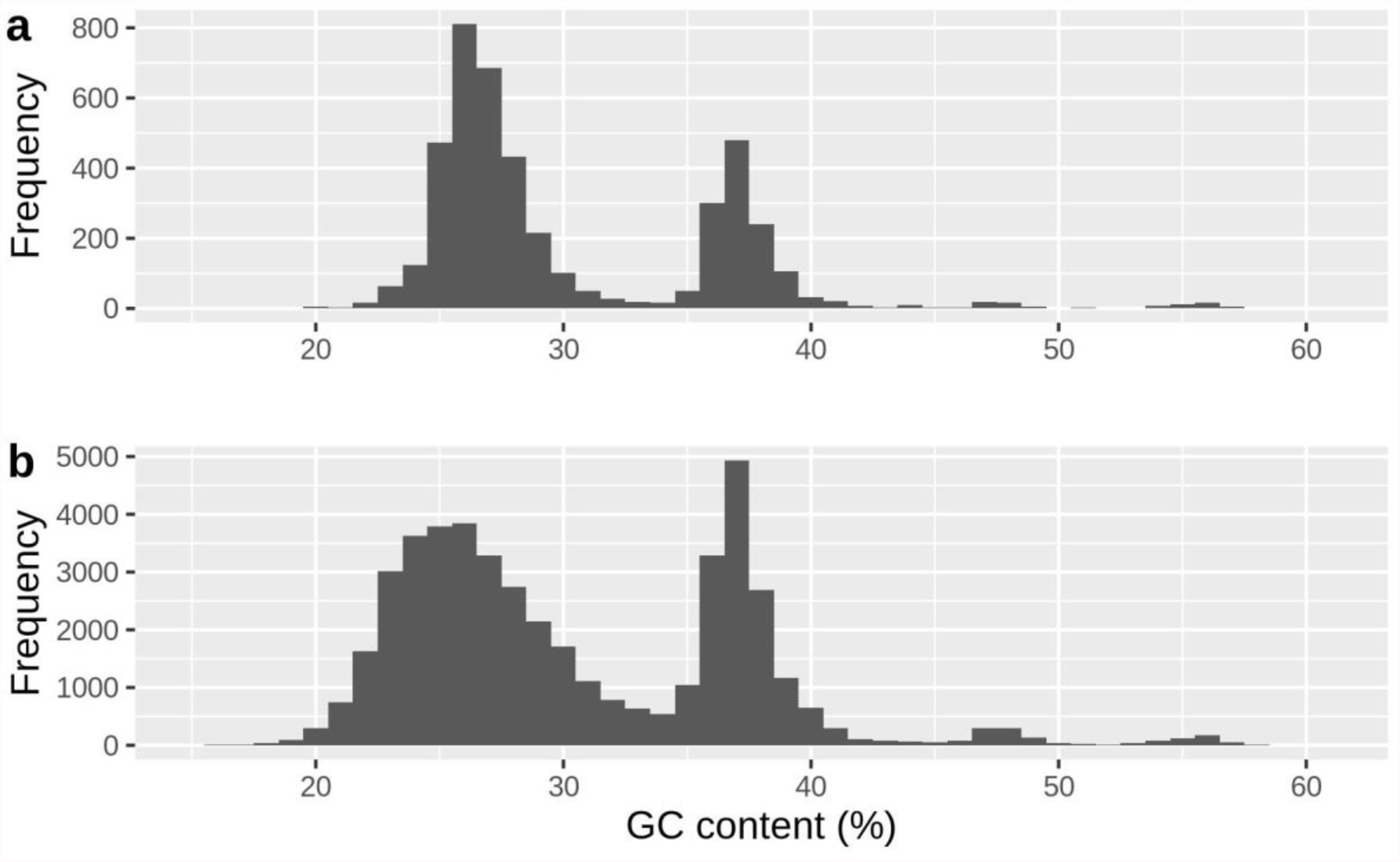
GC-distribution in the reference assembly. **a** distribution based on 50kbp-windows, **b** distribution based on 5kbp-windows.

**Fig. S2:**
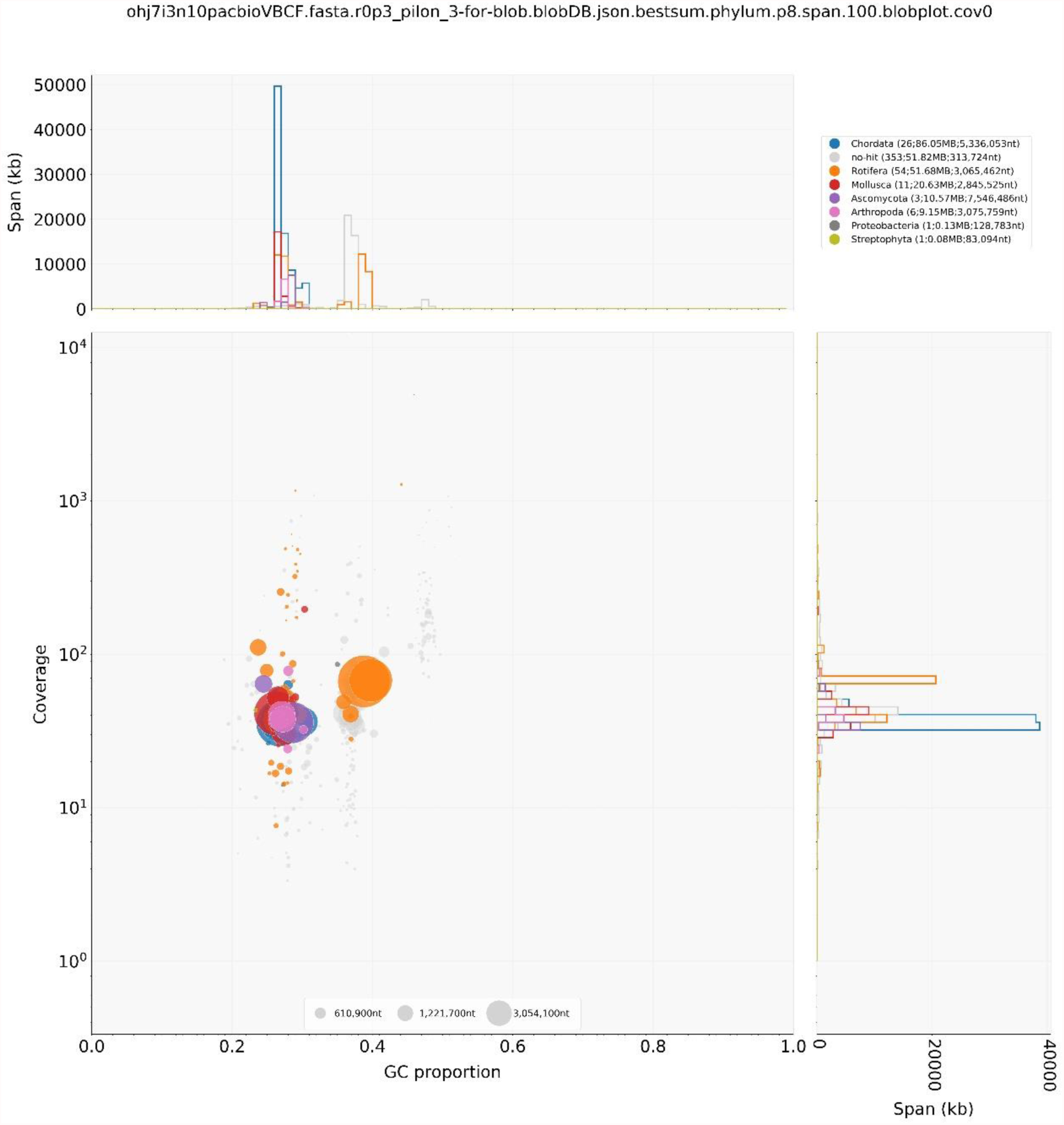
Blobplot of cleaned OHJ7i3n10 reads against reference genome.

**Fig. S3:**
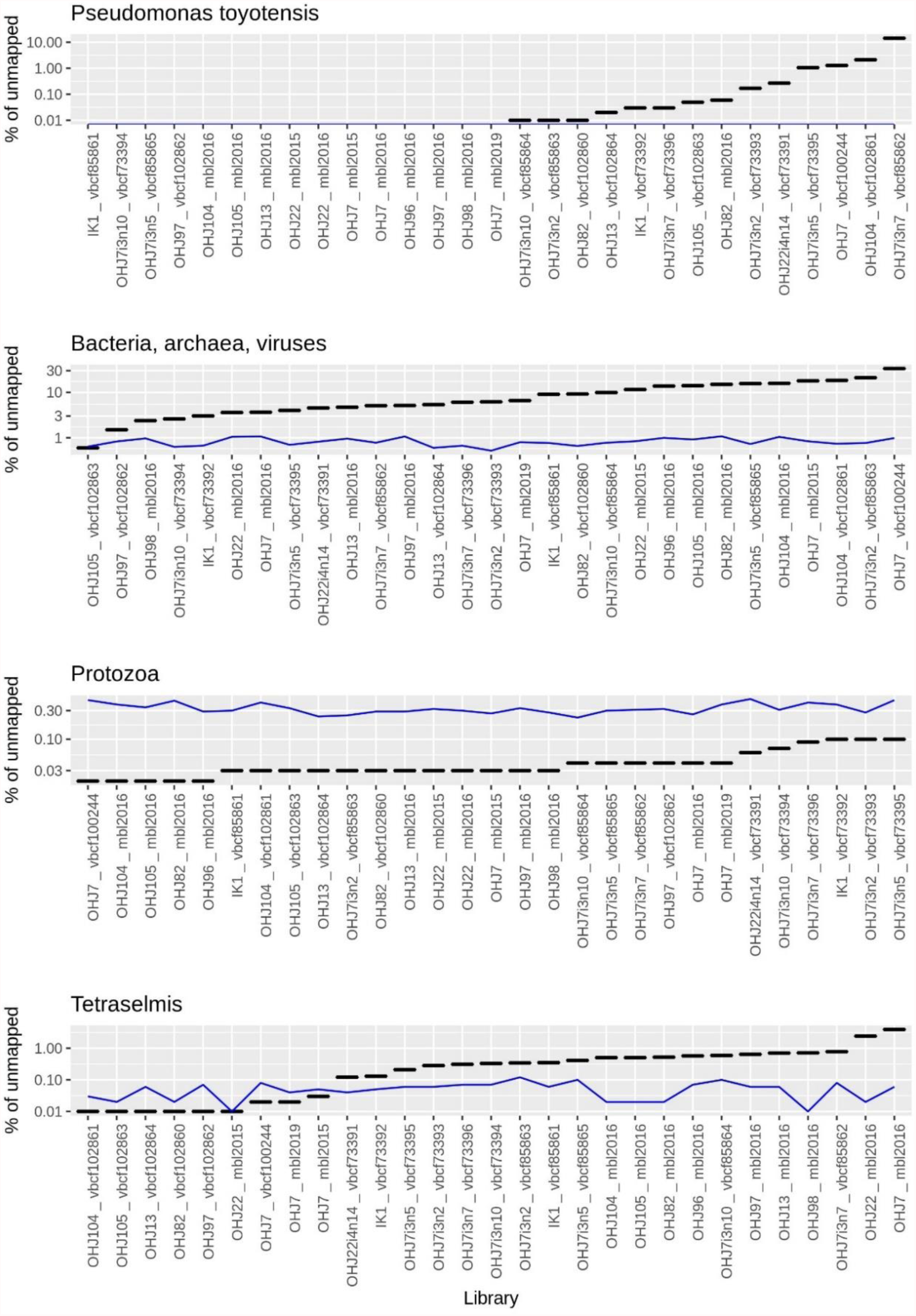
Contamination rates of short-read libraries (before contaminant removal). Horizontal dashes indicate the percentage of reads in the unmapped-reads fraction (i.e., quality-trimmed reads that did *not* align to the reference genome in the first alignment) that could be assigned to four classes of contaminants: *Pseudomonas toyotensis*, (b) other bacteria, archaea, viruses, (c) protozoan contaminants, and (d) *Tetraselmis* (food algae). The blue line indicates the “false-discovery rate”, which was obtained by applying the same pipeline to the mapped-reads fraction of each library.

**Fig. S4:**
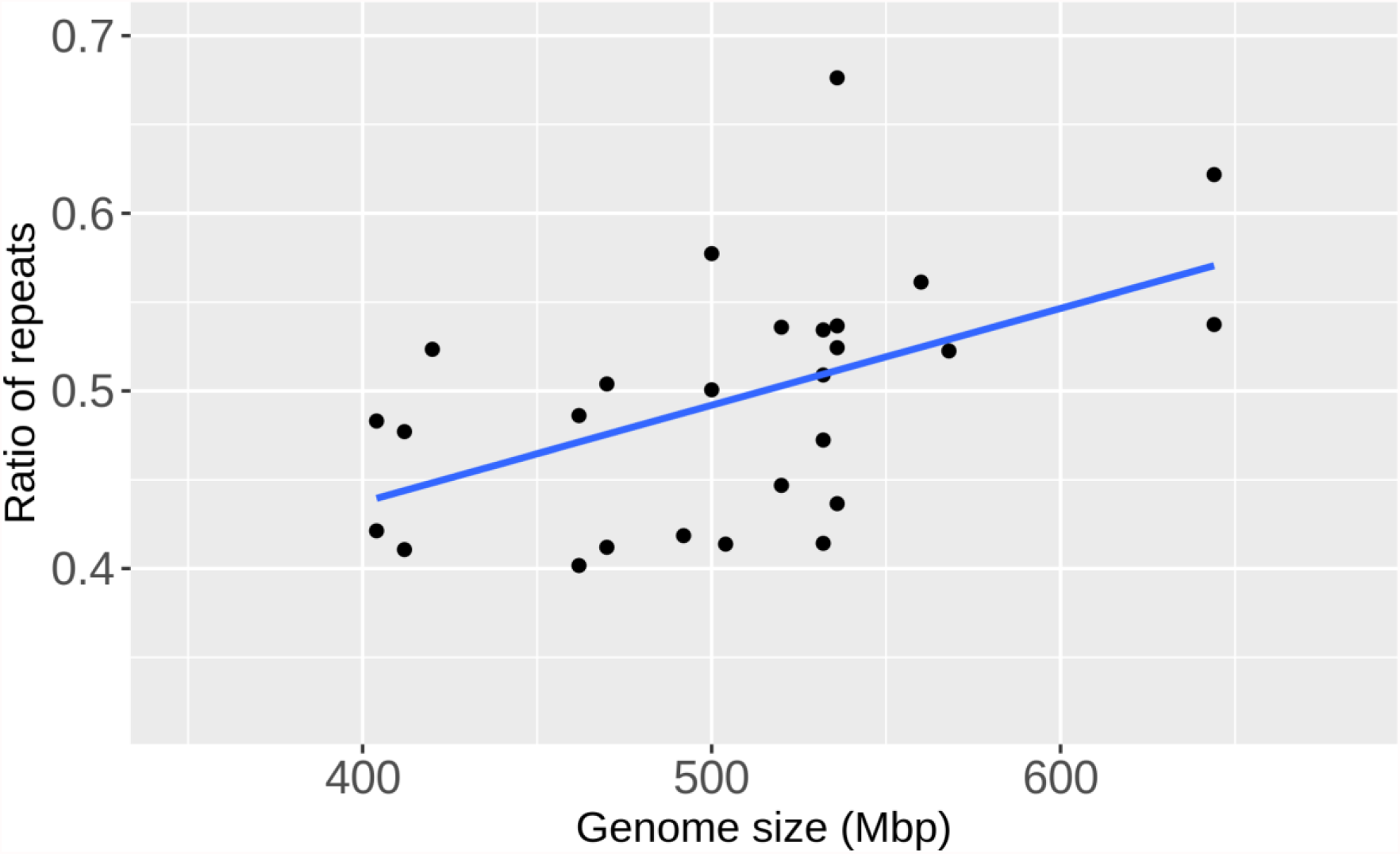
**Genome size (2C, as measured by flow cytometry) versus Ratio of repeats**, a fitted parameter of findGSE [2]. The two variables are significantly correlated with each other (Spearman-rank correlation test, ρ=0.609, *P*=0.0057)

**Fig. S5:**
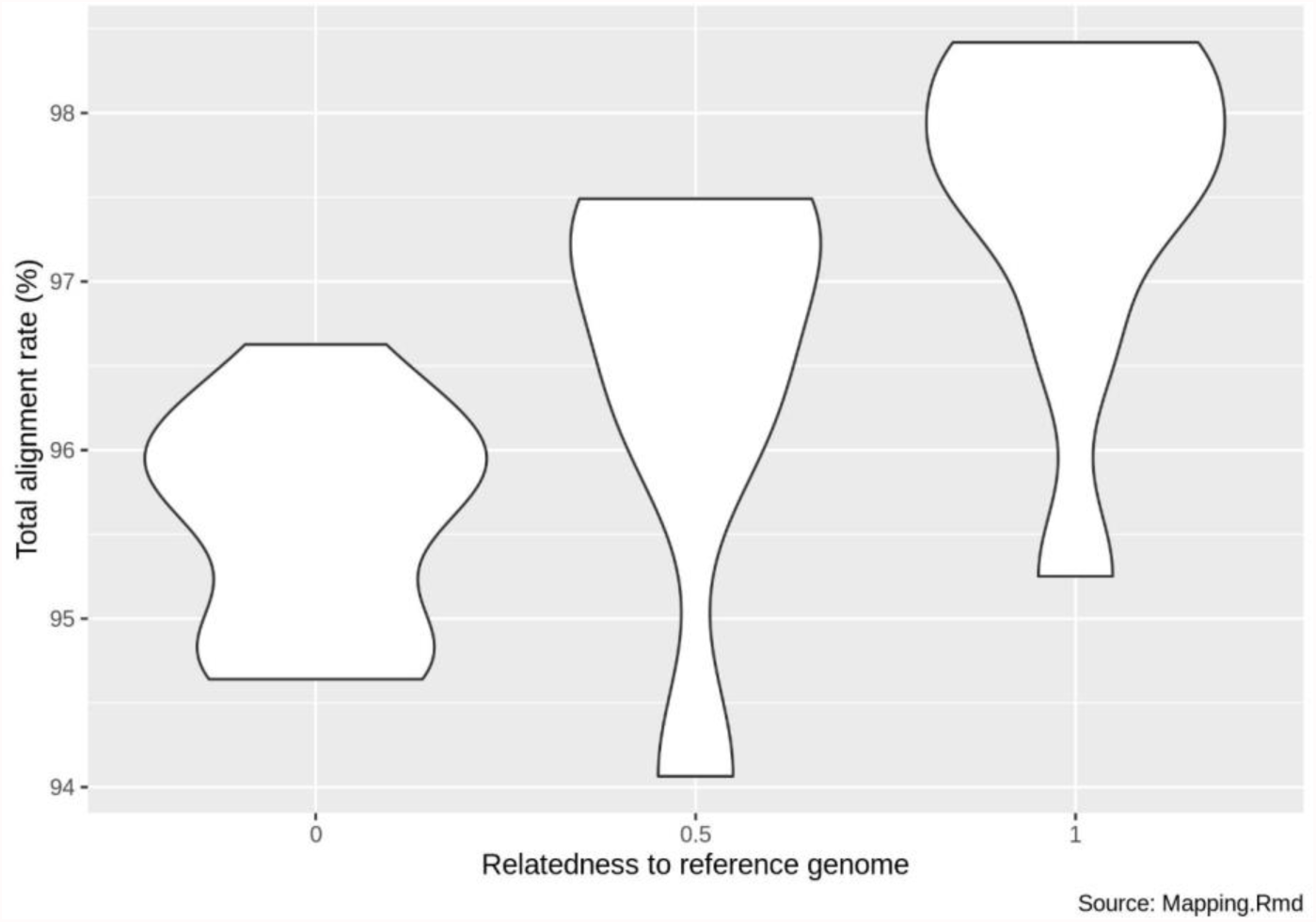
Total alignment rate *vs*. relatedness of DNA source to the reference genome. Relatedness was assumed to be 1 for all clones within the OHJ7i3-selfed line, 0.5 for the inbred line cross (IK1) and the natural ancestor of the inbred line (OHJ7), and 0 for all other clones which hatched from the natural population.

**Fig. S6:**
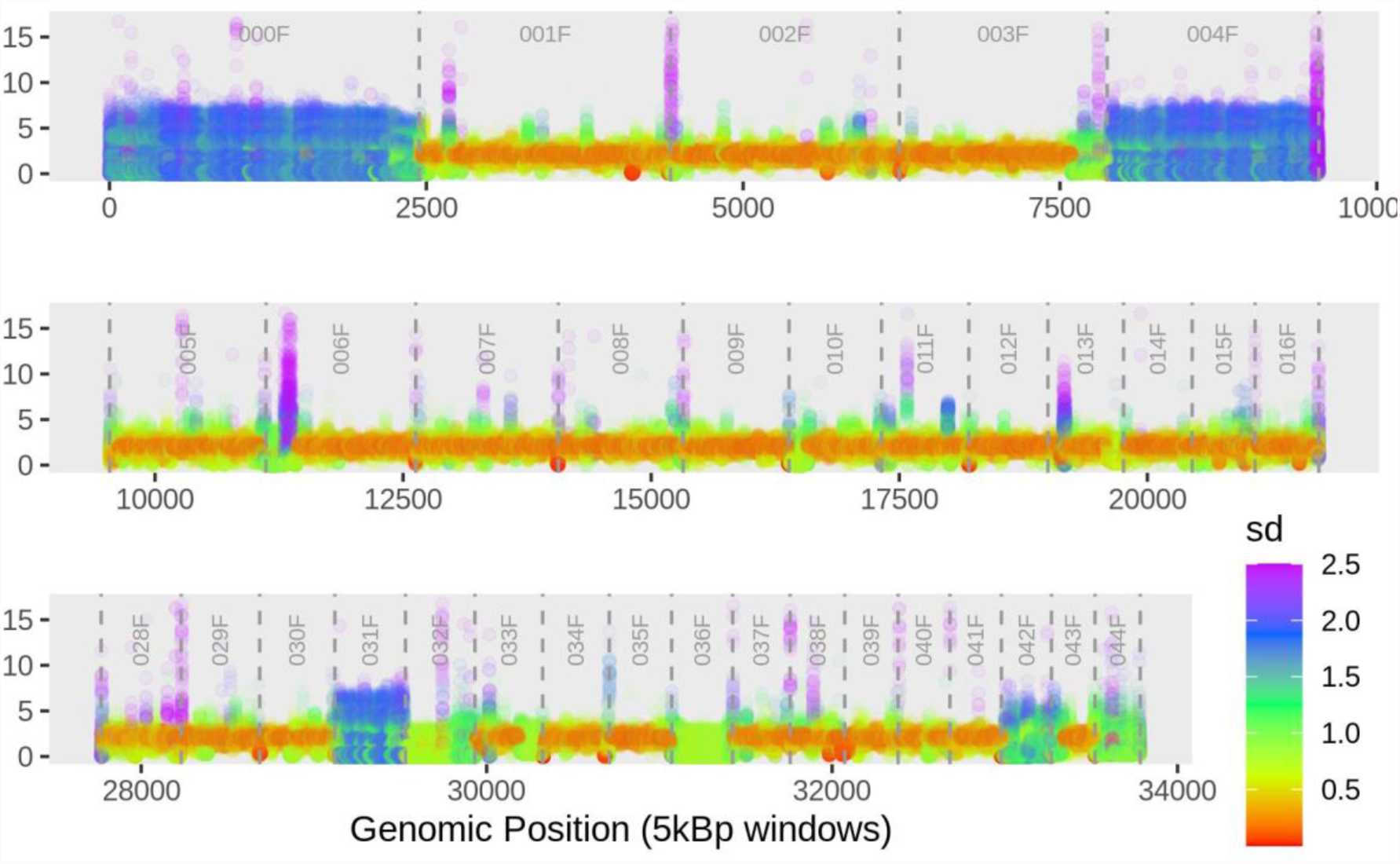
Coverage and coverage variation along representative contigs of the *B. asplanchnoidis* genome (5kBp resolution). Each circle represents a 5kBp window of one of the 31 sequencing libraries. Coverage (y-axis) was normalized by dividing the per-bp coverage of each window by 1/2 of the (mean) exon coverage of the respective library. Contig borders are indicated by vertical dashed lines, and contig IDs are listed on the top. All contigs displayed here span ∼140 Mbp in total, i.e. about 60% of the assembly. Variation across clones/libraries is indicated by color (based on standard deviations). Standard deviations larger than 2.5 (about 4% of the data) had to be capped to a value 2.5, to allow colored visualization.

**Fig. S7:**
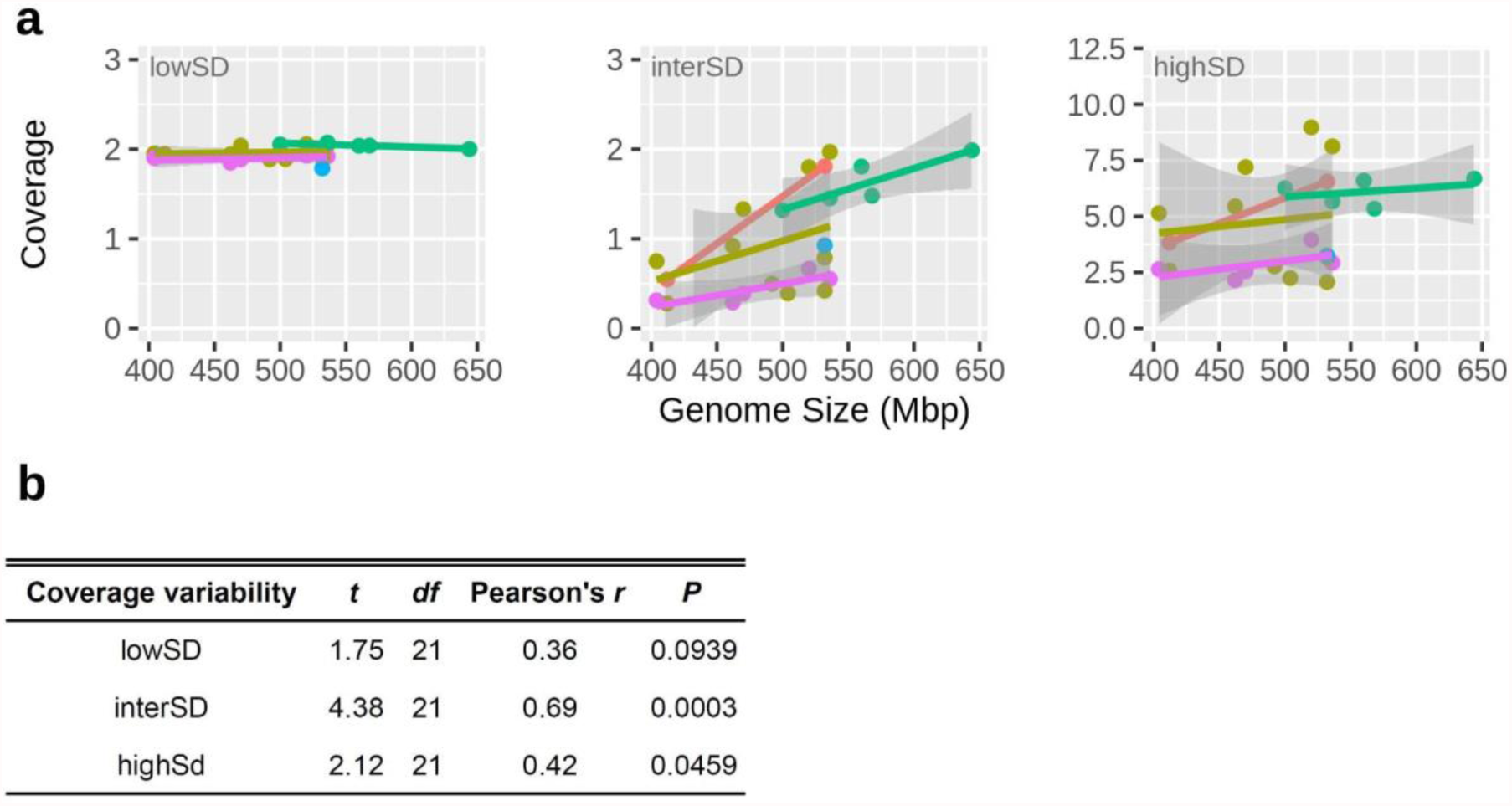
Genomic regions of elevated coverage variability are responsible for genome size variation. This figure shows the same analysis as in Fig. 4 but it excludes the library preps “C” (c.f., Tab. S4), which showed considerably higher coverage and higher GC content in interSD and highSD regions than the other libraries. **a** Mean coverage versus genome size [flow cytometry data on genome size was taken from 3]. Dots represent the mean coverage per library. Colors indicate co-prepared libraries (as in Tab. S4): Orange = A, Pink = B, Green = D, Blue= E, Gold = F. **b** partial correlations between the variables ‘mean coverage’ and ‘genome size’.

**Fig. S8:**
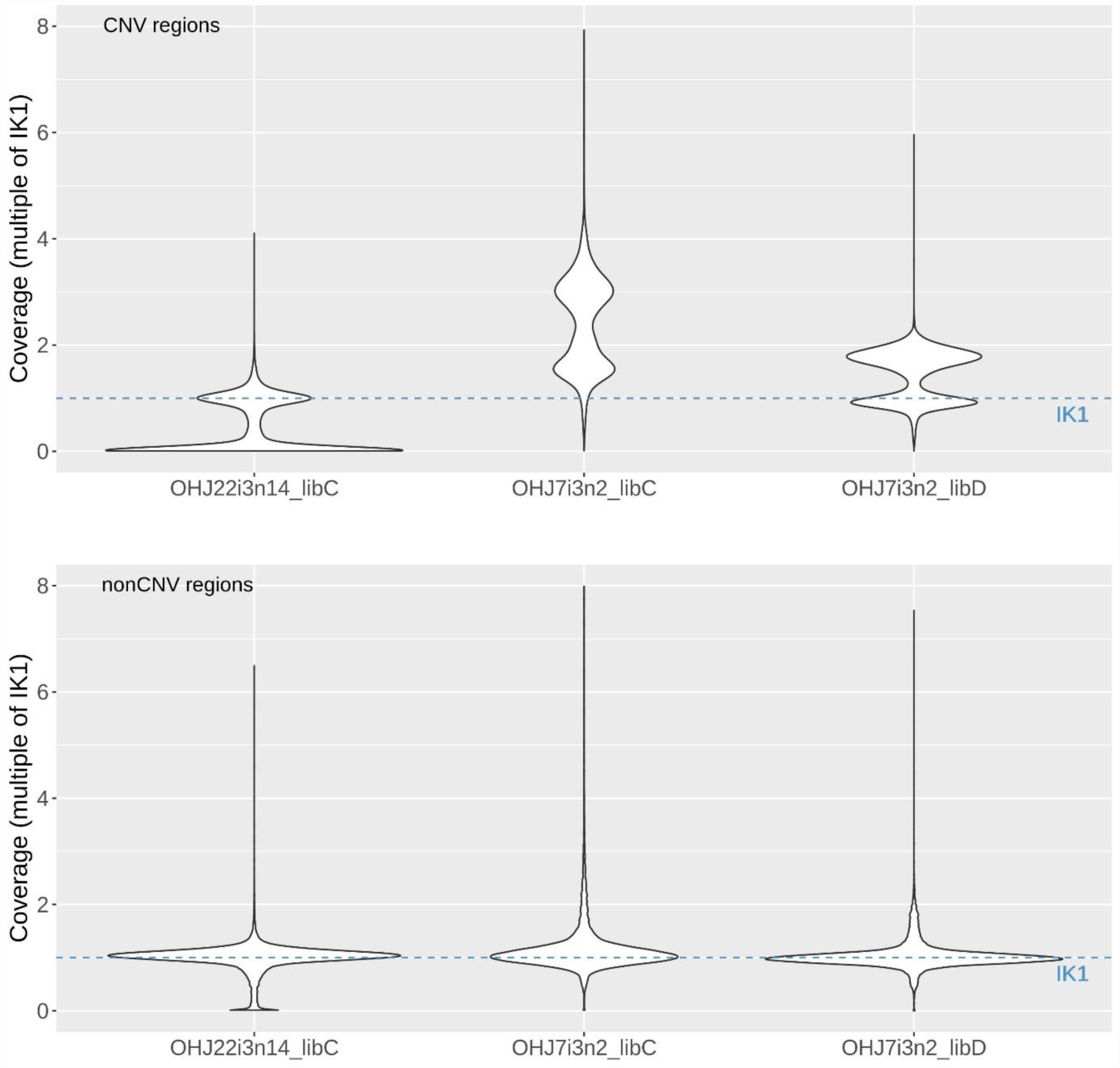
Genome-wide coverage patterns of two parental clones and their cross. This figure summarizes per-base coverage estimates of all 5000bp windows along the genome (subdivided in CNV regions and non-CNV regions, respectively). Coverages of the two parental clones (OHJ22i3n14, OHJ7i3n2) are expressed as multiple of the coverage of the crossed offspring clone (IK1). The dashed line at 1 refers to the crossed offspring IK1. The extensions “libC” and “libD” designate co-prepared libraries as in **Tab S4**.

**Fig. S9:**
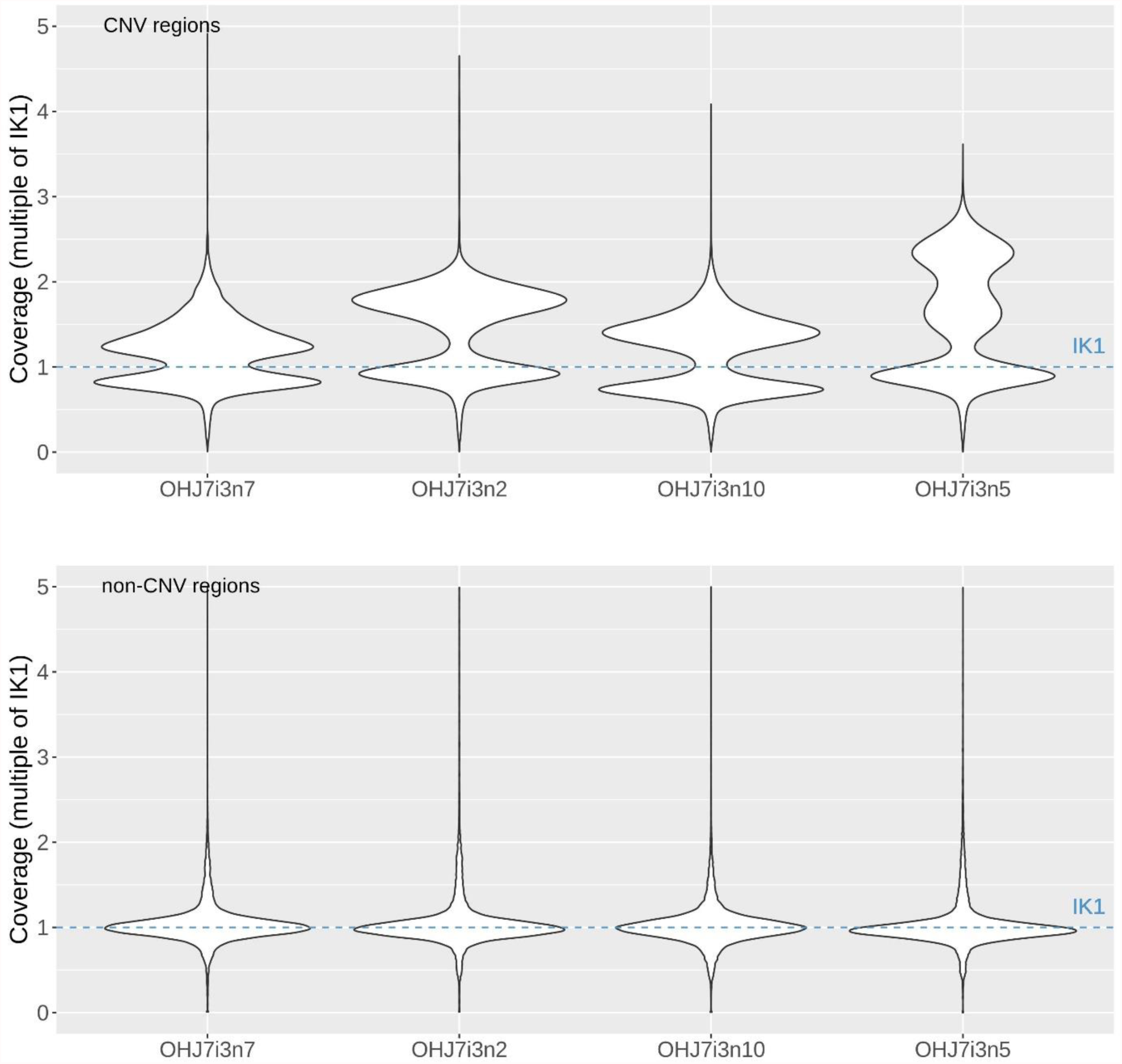
Genome-wide coverage patterns of multiple clones of a selfed line. This figure summarizes per-base coverage estimates of all 5000bp windows along the genome (subdivided in CNV regions and nonCNV regions, respectively). Coverages of the four clones are expressed as multiple of the coverage of the crossed offspring clone (IK1). The dashed line at 1 refers to the crossed offspring IK1. Clones are ordered in ascending genome size: OHJ7i3n7 (536Mbp), OHJ7i3n2 (560MBp), OHJ7i3n10 (568Mbp), OHJ7i3n5 (644Mbp). Only the library preparations “D” Tab. S4 are displayed here, but library preparations “C” exhibited a very similar pattern (not shown).

**Fig. S10:**
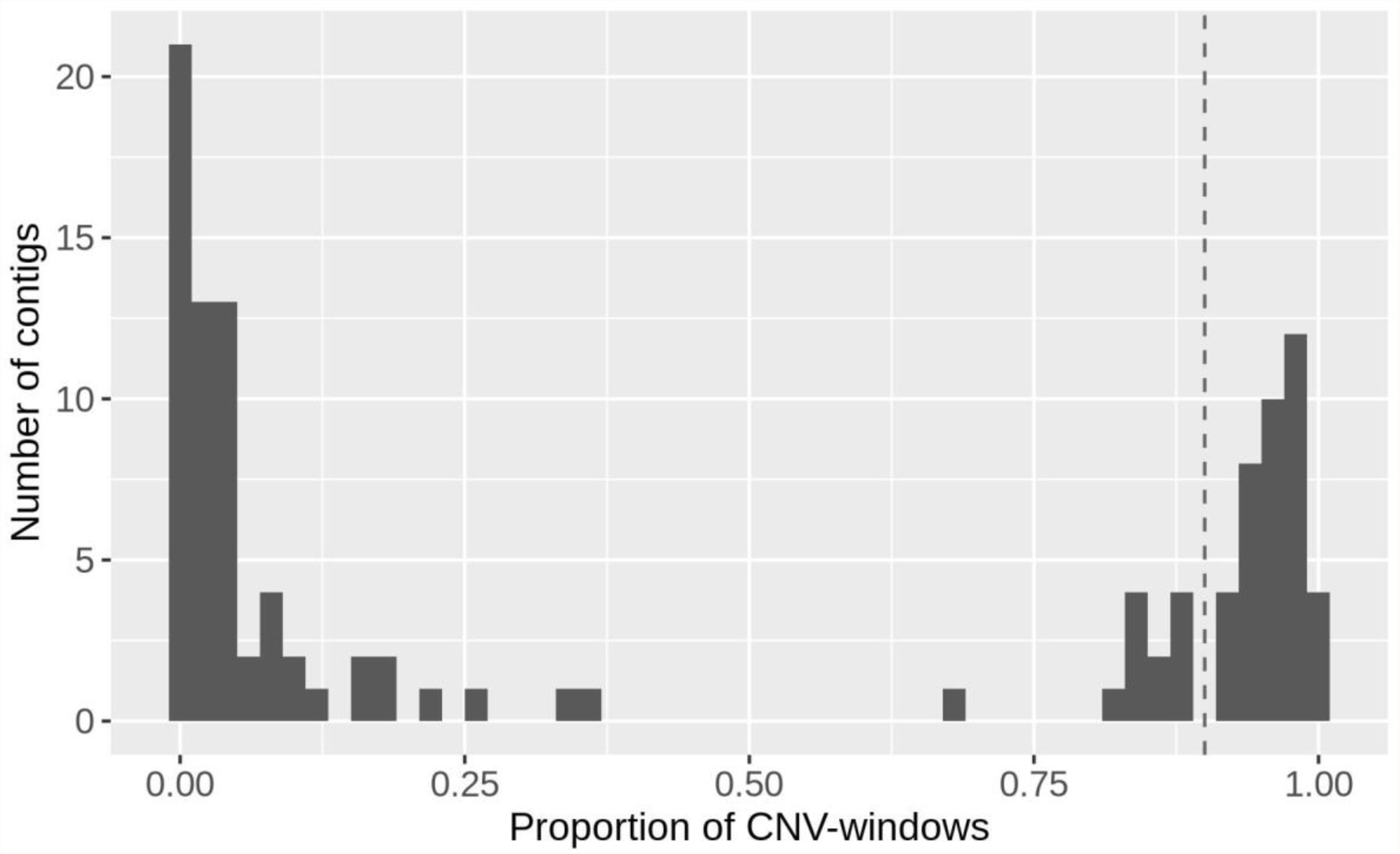
Contig space that is occupied by CNVs. The X-axis displays the proportion of all 5kbp windows in a contig that have been classified as part of a CNV. For this graph, we only used contigs with at least 50 windows in length (i.e., contig length > 250kbp; n=114). The dashed line is our threshold at 0.9 for defining ‘B-contigs’.

**Fig. S11:**
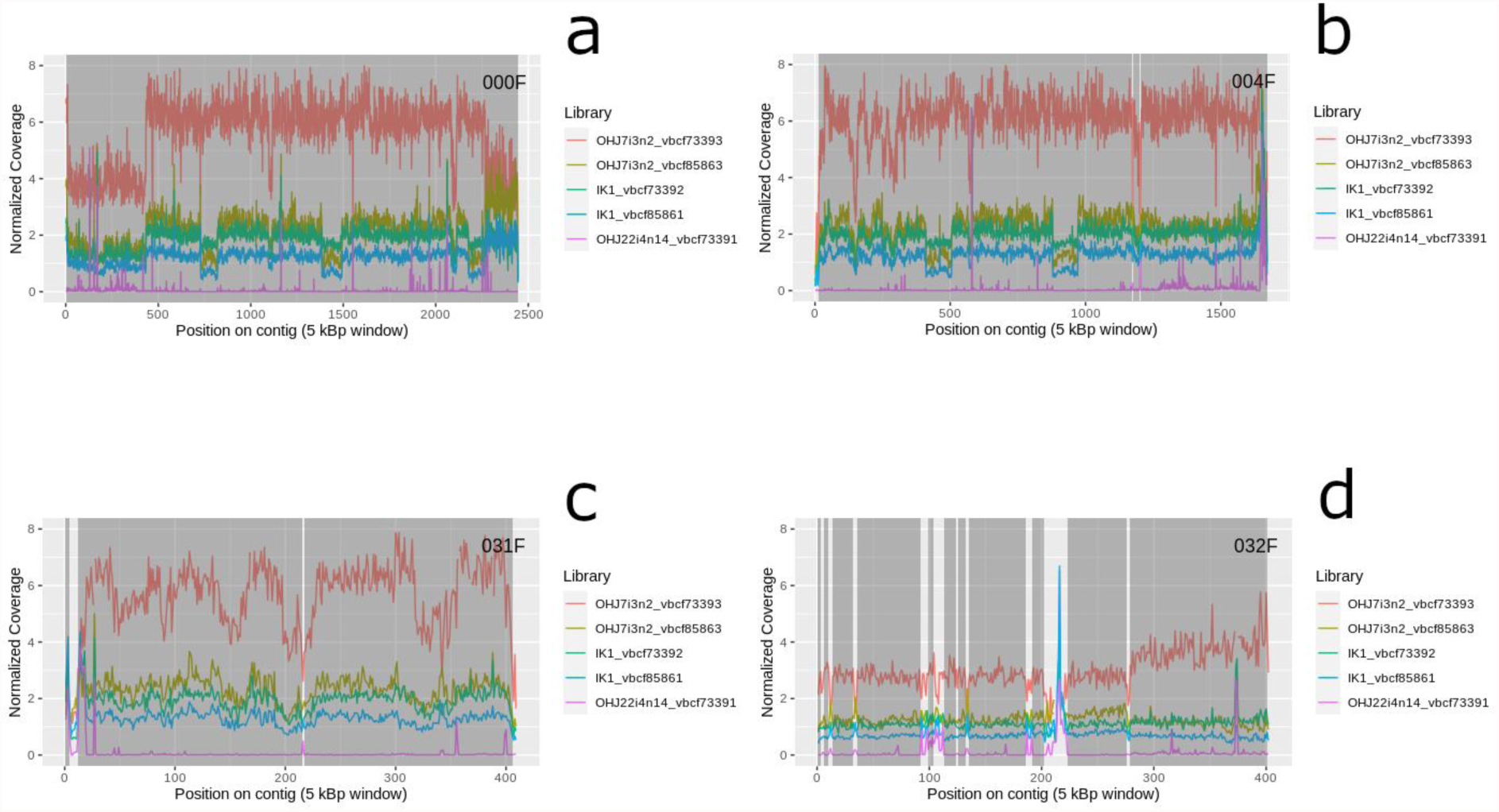
Examples of “B-contigs”. B-contigs consist almost exclusively of coverage-variable regions, with a consistent coverage pattern. Shaded regions highlight calls of individual CNVs, while open regions are coverage breakpoints, either due to low coverage variation and/or shifts in the coverage patterns among clones. Colored lines represent normalized coverage values of different rotifer clones/libraries (with “2” equaling the coverage at exonic regions). The displayed clones are a “trio” consisting of the parents (OHJ7i3n2, 2C genome size: 560 Mbp; OHJ22i4n14, 420 Mbp) and their sexual offspring (IK1, 500 Mbp).

**Fig. S12:**
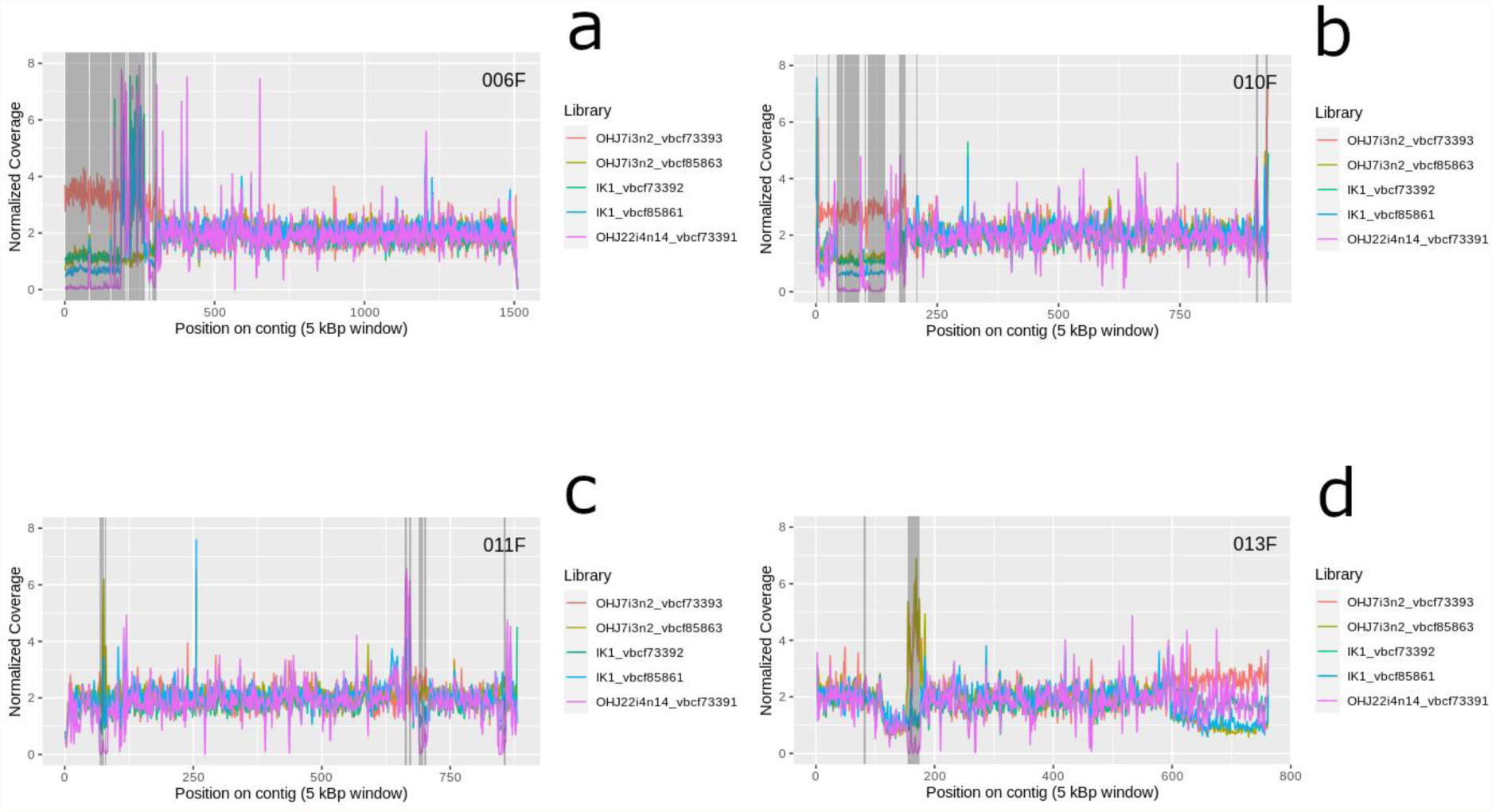
Example of small-scale coverage variation. The contigs displayed here contain shorter sections that have been identified as CNVs. Shaded regions highlight calls of individual CNVs, while open regions are coverage breakpoints, either due to low coverage variation and/or shifts in the coverage patterns among clones. Colored lines represent normalized coverage values of different rotifer clones/libraries (with “2” equaling the coverage at exonic regions). The displayed clones are a “trio” consisting of the parents (OHJ7i3n2, 2C genome size: 560 Mbp; OHJ22i4n14, 420 Mbp) and their sexual offspring (IK1, 500 Mbp).

**Fig. S13:**
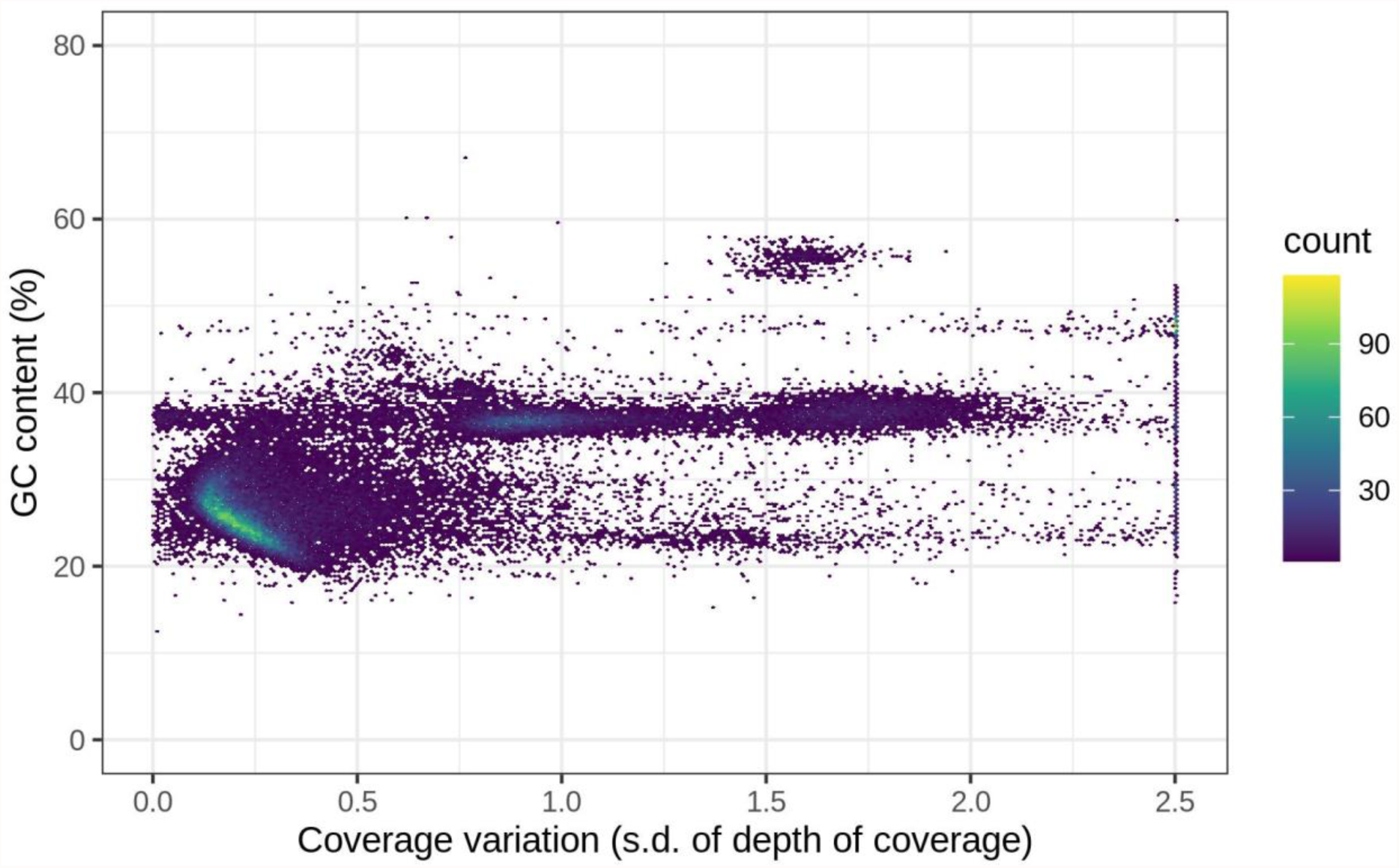
GC-content vs. coverage variability in different regions of the genome of *B. asplanchnoidis*. Hexbin chart is based on all 5kbp windows of the assembly (bins=250, in ggplot2).

**Fig. S14:**
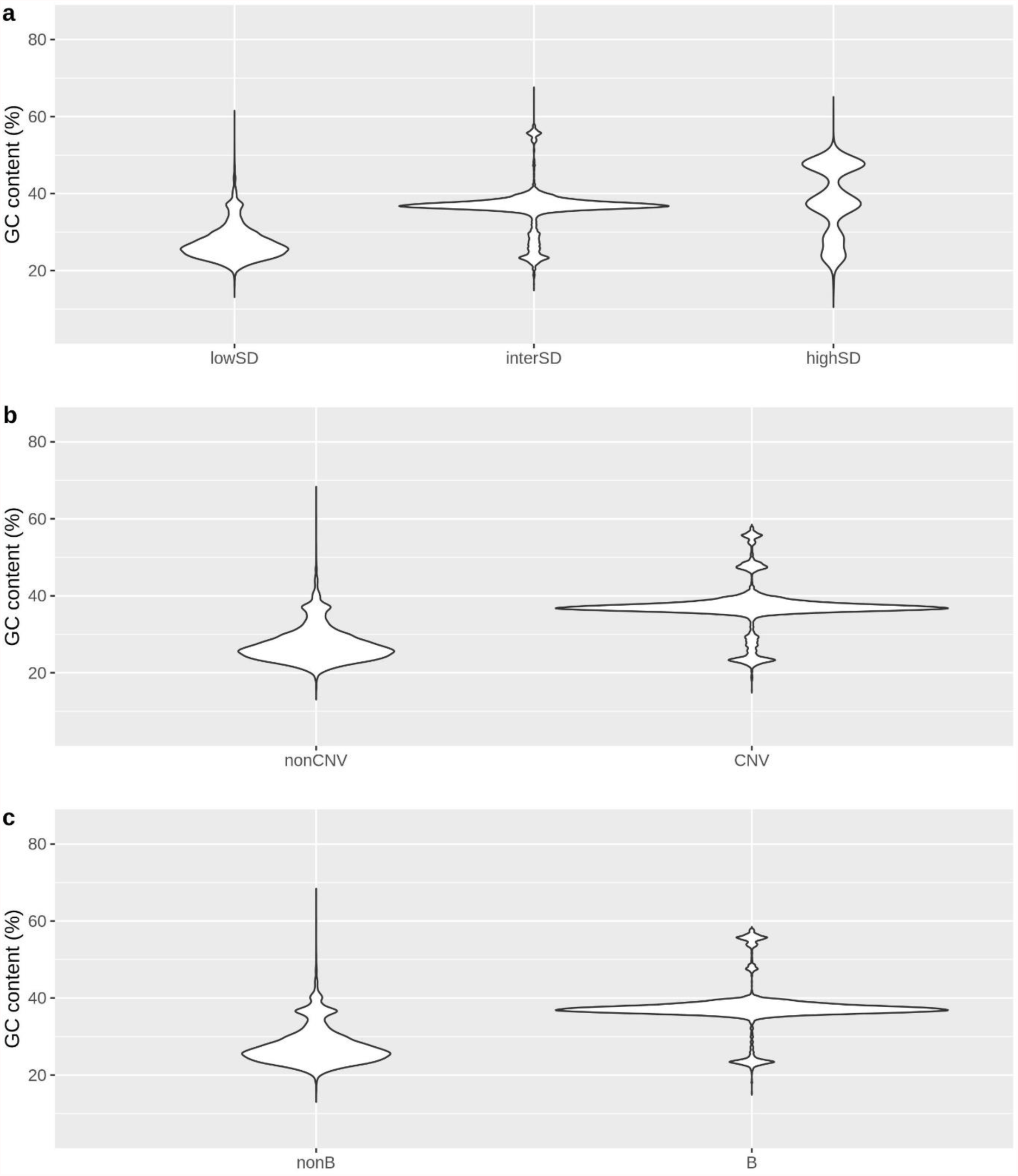
GC-content of genomic regions for in the *B. asplanchnoidis* genome. a. Regions defined by coverage variability, **b** CNV regions (i.e., elevated coverage variability & locally consistent coverage patterns), **c** B-contigs (contigs consisting of >90% CNVs). For **c**, only contigs with at least 50 windows were considered. The distributions shown here are based on GC-contents calculated from 5000-bp windows. To prevent losses in precision, the last window of a contig was excluded (since it was always shorter than 5000 bp).

**Fig. S15:**
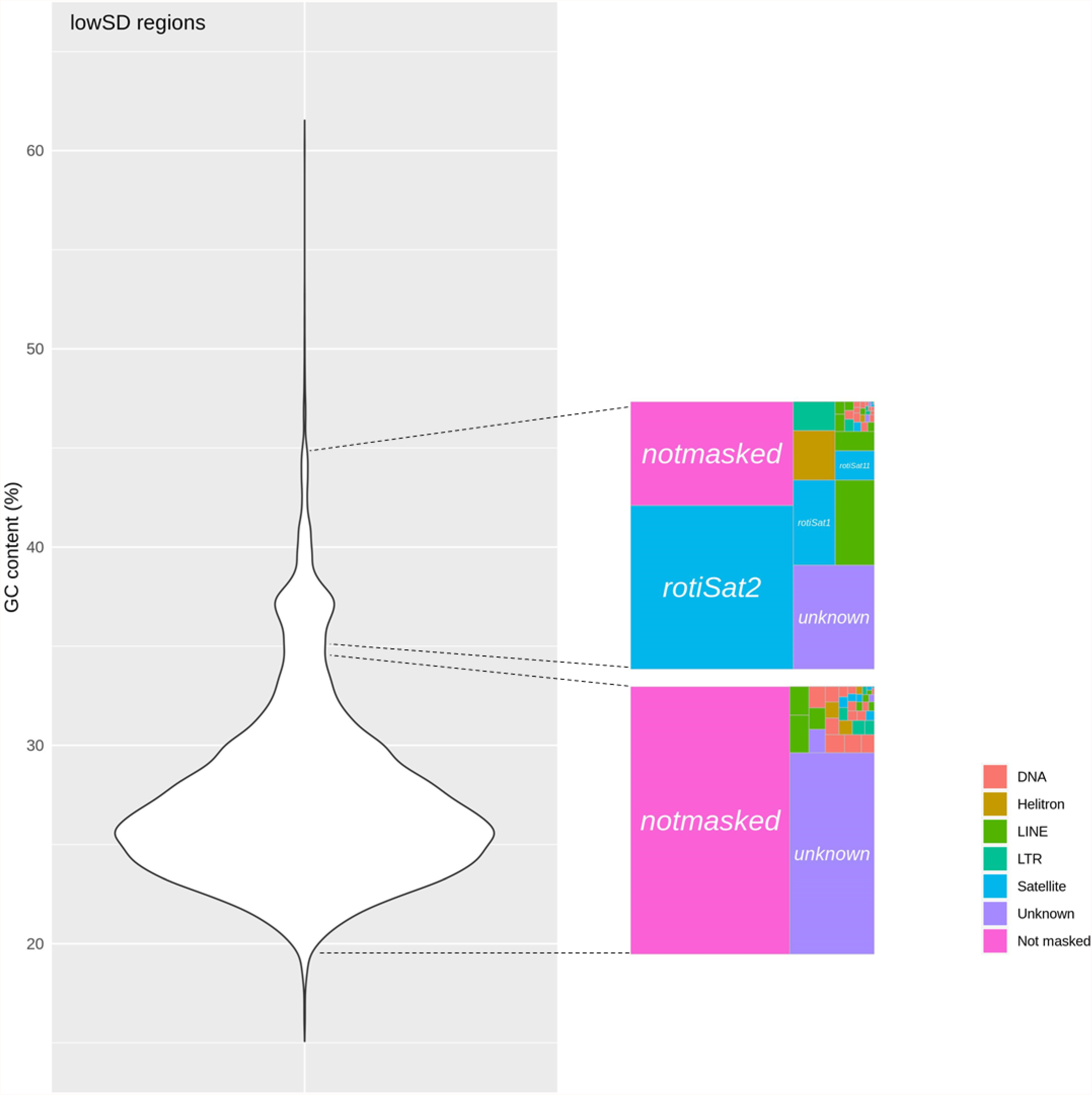
Repeat composition of various GC fractions of lowSD regions. ‘Unknown’ refers to regions that were masked as repeats but could not be ascribed to any of the above categories. ‘Not masked’ refers to regions that were not identified as repeats by repeatModeler2.

**Fig. S16:**
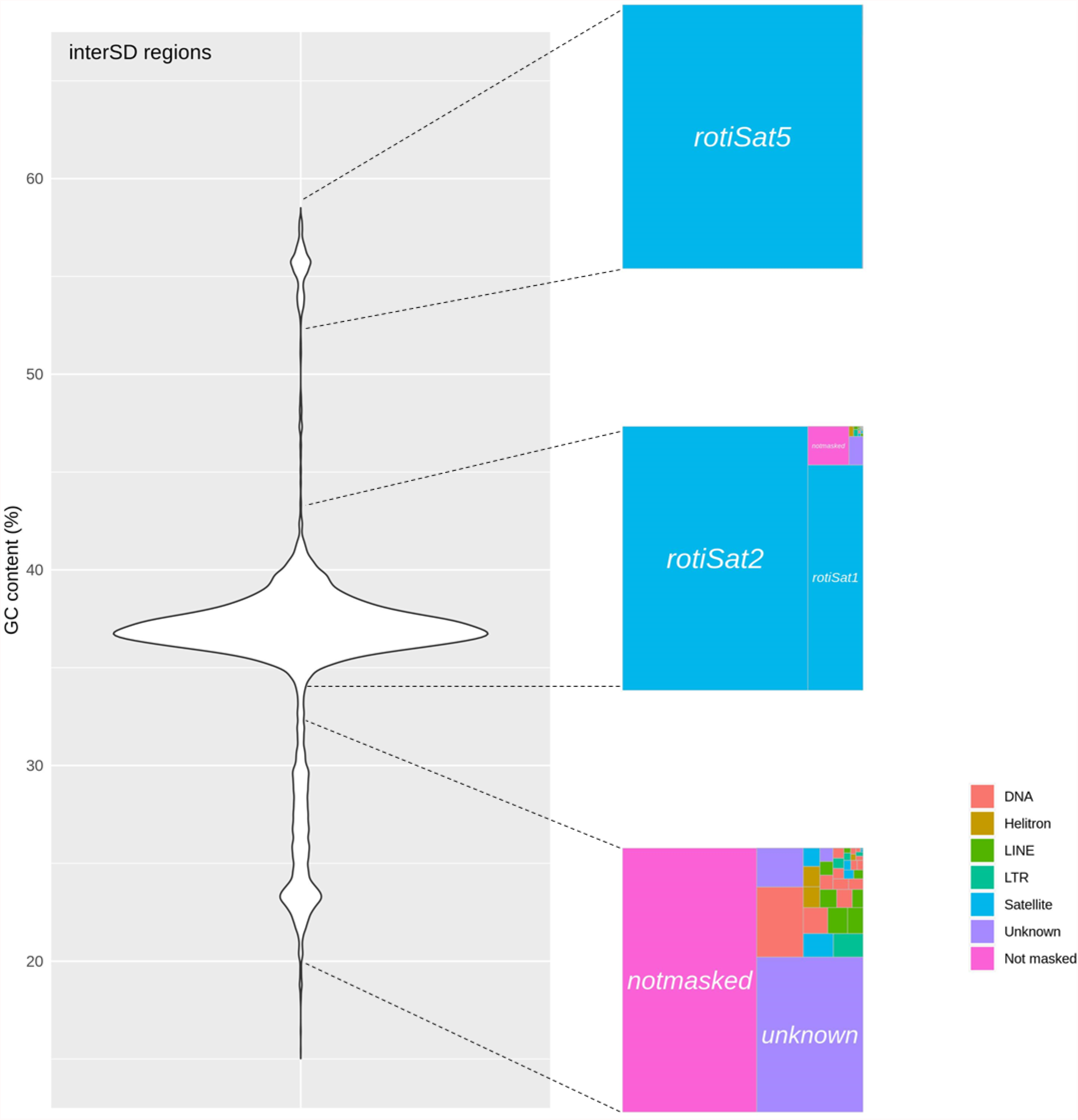
Repeat composition of various GC fractions of interSD regions.

**Fig. S17:**
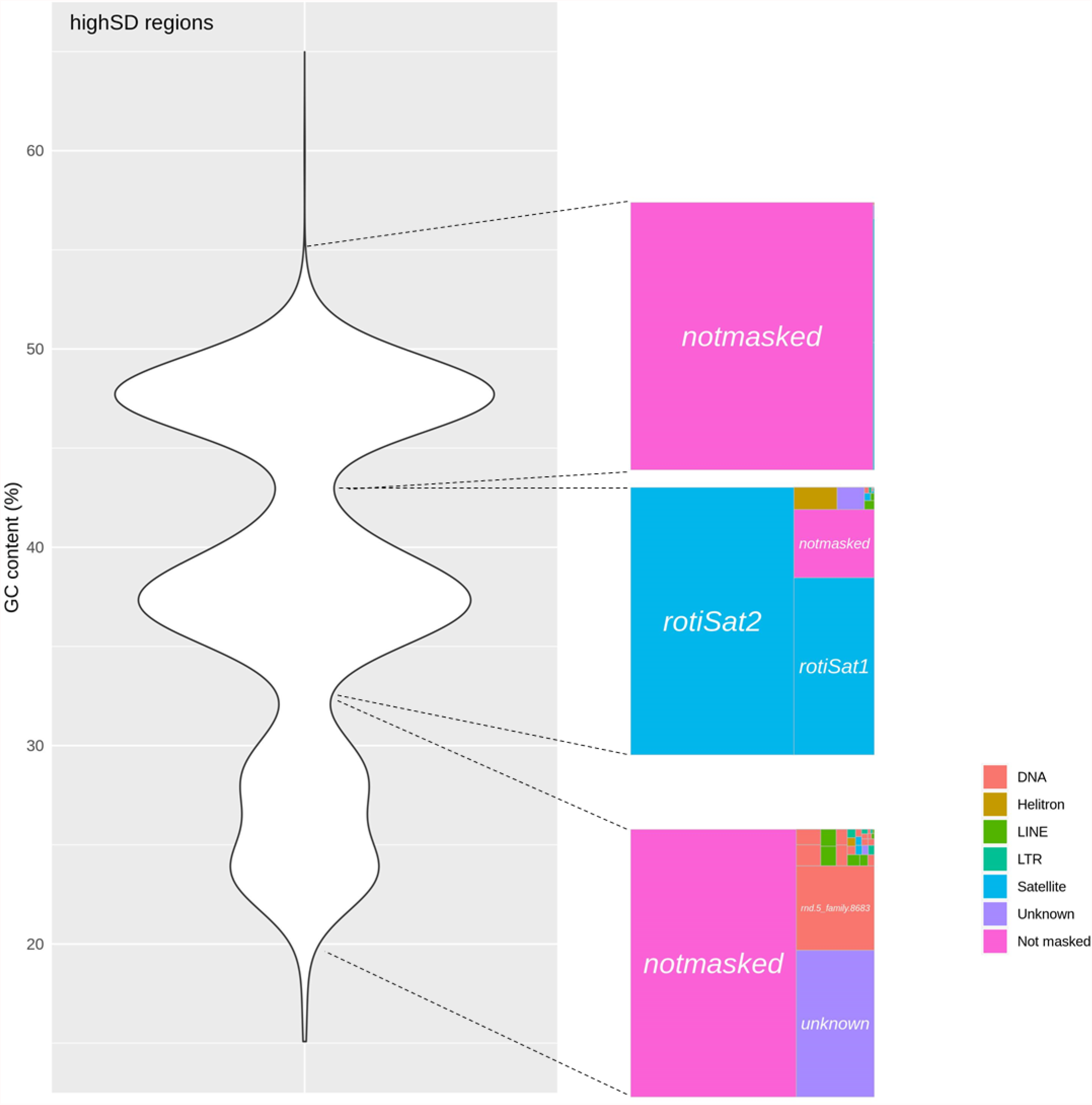
Repeat composition of various GC fractions of highSD regions.

**Fig. S18:**
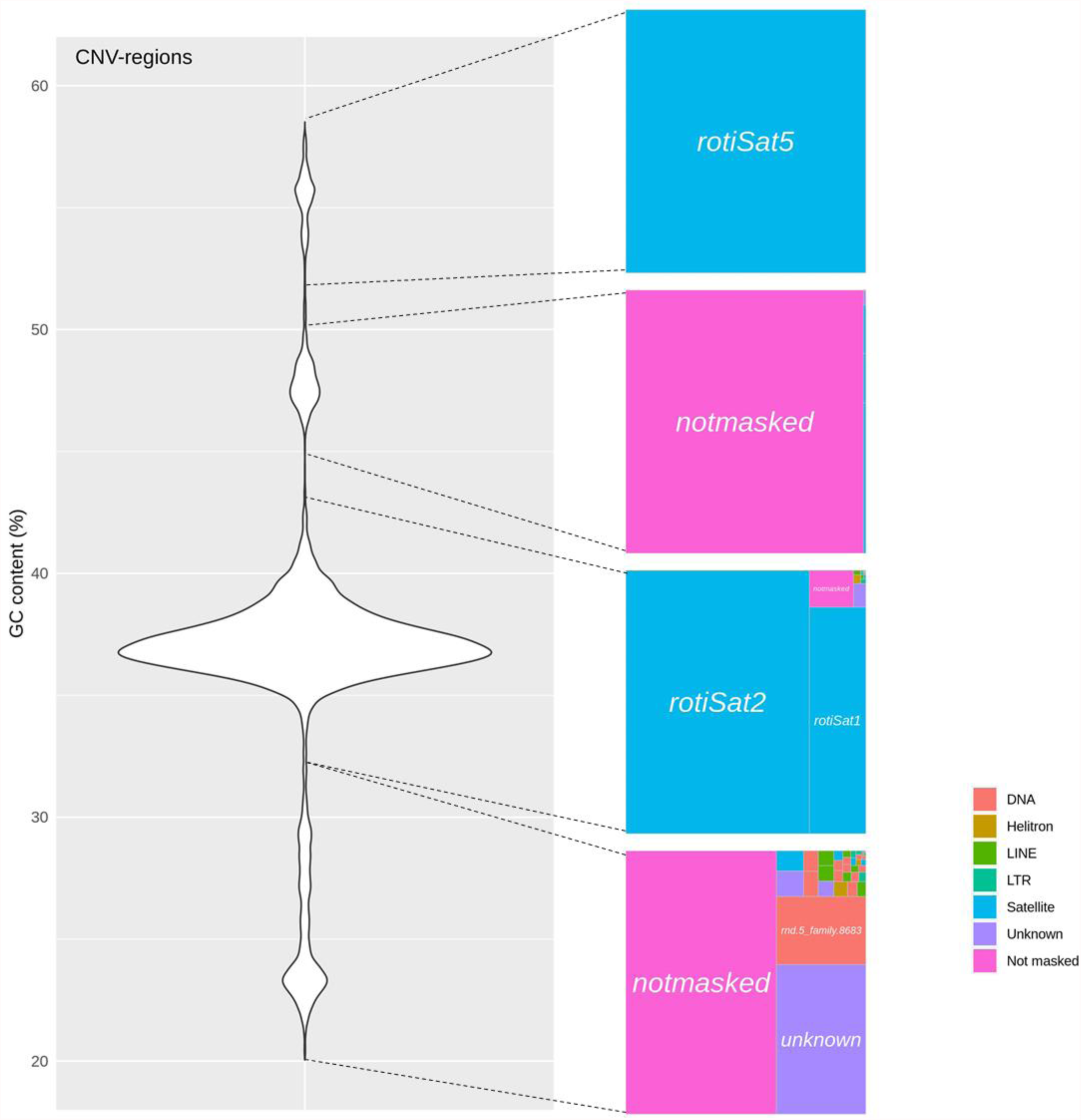
Repeat composition of various GC fractions of CNV regions.

**Fig. S19:**
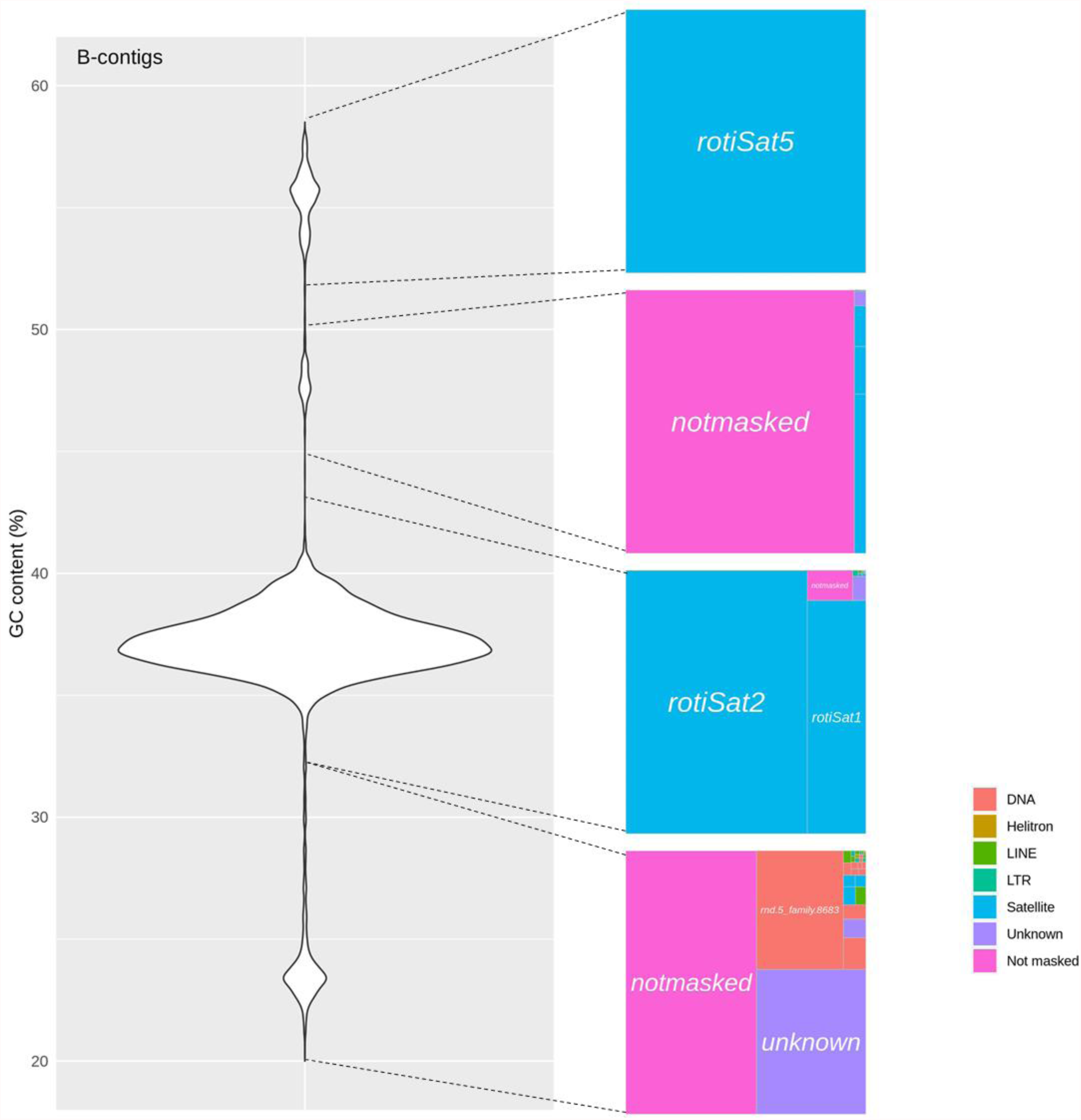
Repeat composition of various GC fractions of B-contigs.

**Table S1:**
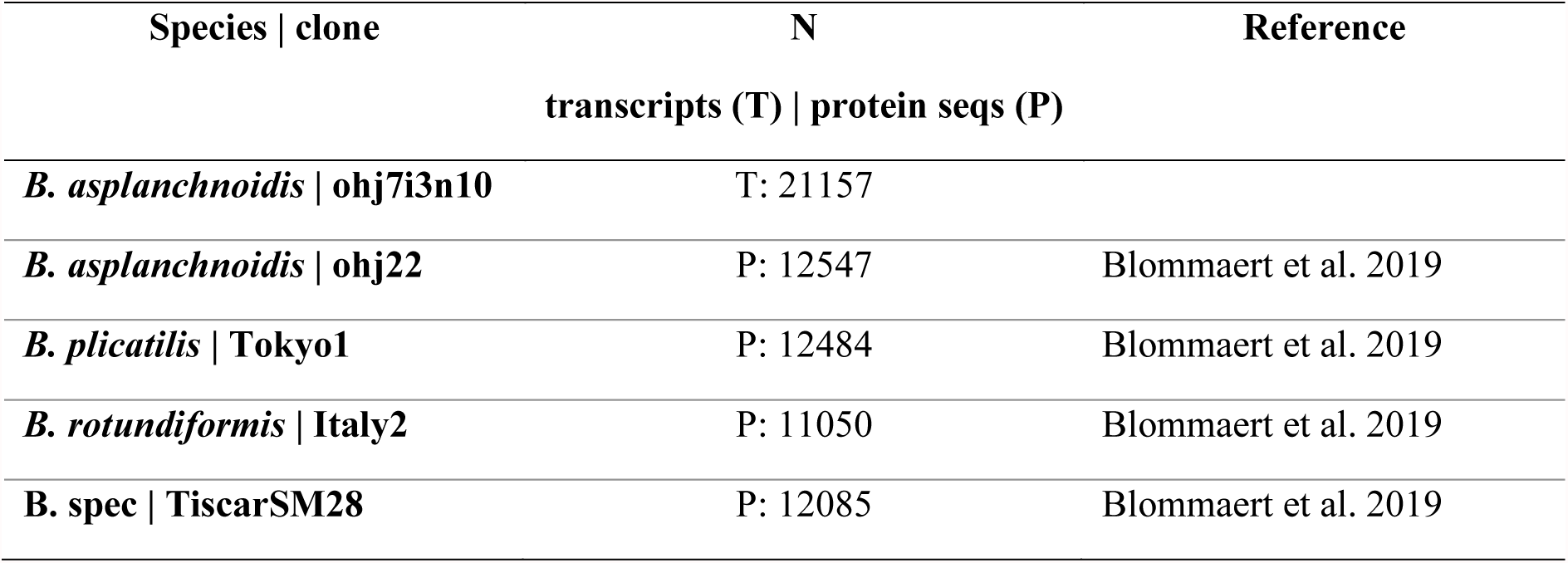
Outline of transcriptome and proteome libraries used for gene annotation.

**Table S2:** **BUSCO completeness statistics** of *B. asplanchniodis* genome in comparison to two previously published genomes of related *Brachionus* species.

| BUSCO (Metazoa n=978) | *B. asplanchnoidis* | *B. plicatilis* | *B. calyciflorus* |
| --- | --- | --- | --- |
| Complete % | 90.6 | 95.0 | 90.7 |
| Single complete % | 87.9 | 91.7 | 88.0 |
| Duplicated complete % | 2.7 | 3.3 | 2.7 |
| Fragmented % | 2.0 | 2.0 | 2.0 |
| Missing % | 7.4 | 3.0 | 7.3 |

Data sources: Published genome of *B. plicatilis* by [4] and of *B. calyciflorus* by [5]

**Table S3:**
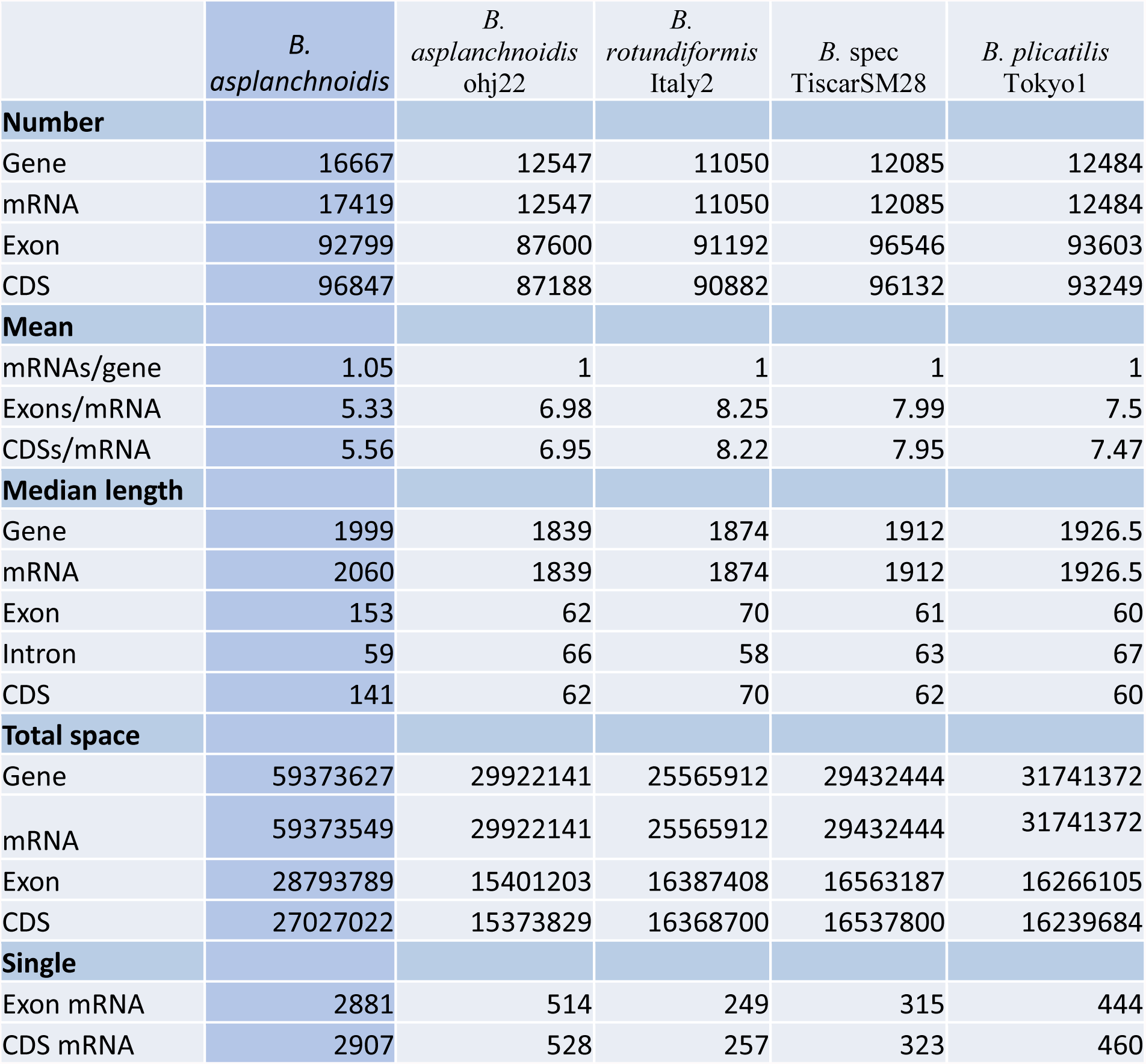
**Summary statistics of B. *asplanchnoidis* annotation** in comparison to annotations of previously published genomes from related *Brachionus* species [6]. For reasons of data availability, this comparison does not include more species of the *B. plicatilis* species complex.

**Table S4:** Information on short-read sequencing libraries.

| Clone | Origin | GS<sup>1</sup><br>(Mbp) | Method | Prep.<sup>2</sup> | max. FS<sup>3</sup> (Bp) | avg. FS<sup>3</sup> (Bp) |
| --- | --- | --- | --- | --- | --- | --- |
| OHJ82 | natural clone | 404 | westburg | F | 2000 | 503 |
|  |  |  | kapa | B | 600 | 450 |
| OHJ22 | natural clone | 412 | kapa | A | 600 | 450 |
|  |  |  | kapa | B | 600 | 450 |
| OHJ104 | natural clone | 462 | westburg | F | 2000 | 499 |
|  |  |  | kapa | B | 600 | 450 |
| OHJ97 | natural clone | 470 | westburg | F | 2000 | 499 |
|  |  |  | kapa | B | 600 | 450 |
| OHJ96 | natural clone | 492 | kapa | B | 600 | 450 |
| OHJ98 | natural clone | 504 | kapa | B | 600 | 450 |
| OHJ105 | natural clone | 520 | westburg | F | 2000 | 493 |
|  |  |  | kapa | B | 600 | 450 |
| OHJ7 | natural clone | 532 | westburg | E | 1000 | 386 |
|  |  |  | kapa | A | 600 | 450 |
|  |  |  | kapa | B<sup>4</sup> | 600 | 450 |
|  |  |  | kapa | B<sup>4</sup> | 600 | 450 |
| OHJ13 | natural clone | 536 | westburg | F | 2000 | 516 |
|  |  |  | kapa | B | 600 | 450 |
| OHJ22i3n14 | selfed line | 420 | westburg | C | 1000 | 545 |
| OHJ7i3n7 | selfed line | 536 | westburg | C | 1000 | 495 |
|  |  |  | westburg | D | 3000 | 1149 |
| OHJ7i3n2 | selfed line | 560 | westburg | C | 1000 | 632 |
|  |  |  | westburg | D | 3000 | 1228 |
| OHJ7i3n10 | selfed line | 568 | westburg | C | 1000 | 589 |
|  |  |  | westburg | D | 3000 | 1093 |
| OHJ7i3n5 | selfed line | 644 | westburg | C | 1000 | 496 |
|  |  |  | westburg | D | 3000 | 1129 |
| IK1 | selfed line cross | 500 | westburg | C | 1000 | 514 |
|  |  |  | westburg | D | 2000 | 1001 |

<sup>1</sup> genome size (2C) estimated by flow cytometry in [3]
<sup>2</sup> Indicates co-prepared libraries
<sup>3</sup> Fragment size (FS) of the library
<sup>4</sup> Both derive from the same original preparation, but one was re-sequenced and had an intermediate PCR-step

**Table S5:**
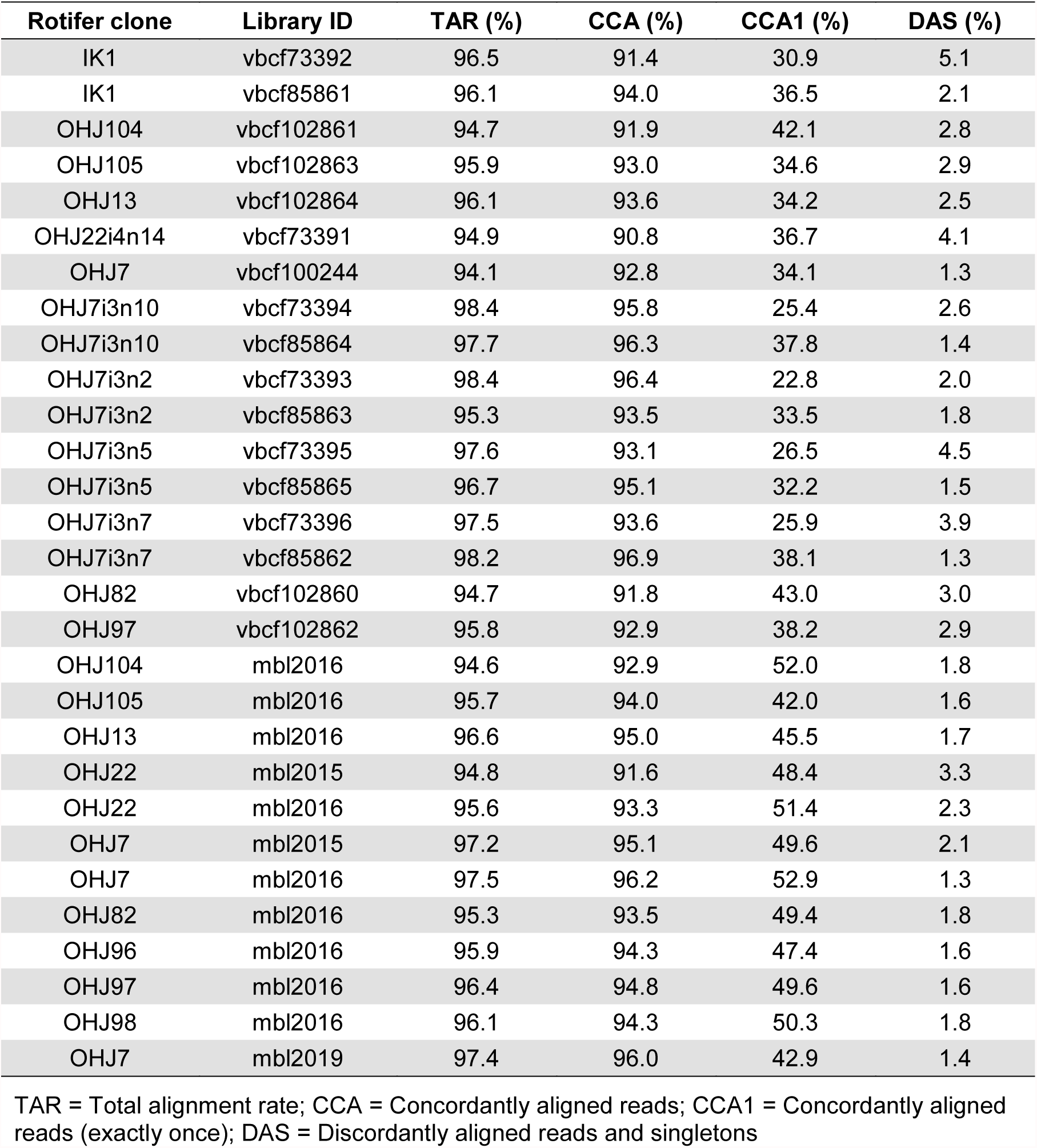
Alignment statistics of short reads to reference genome.

**Table S6:** Classification of ‘B-contigs’ in terms of the proportion of 5kbp windows that are CNV.

| **Contig** | **Total 5kbp windows** | **CNV-windows** | **Proportion** |
| --- | --- | --- | --- |
| 000F | 2444 | 2436 | 0.997 |
| 057F | 166 | 165 | 0.994 |
| 058F | 163 | 162 | 0.994 |
| 071F | 109 | 108 | 0.991 |
| 004F | 1671 | 1651 | 0.988 |
| 083F | 79 | 78 | 0.987 |
| 084F | 78 | 77 | 0.987 |
| 086F | 72 | 71 | 0.986 |
| 048F | 210 | 207 | 0.986 |
| 091F | 66 | 65 | 0.985 |
| 099F | 59 | 58 | 0.983 |
| 059F | 155 | 152 | 0.981 |
| 109F | 50 | 49 | 0.980 |
| 061F | 146 | 143 | 0.979 |
| 055F | 175 | 171 | 0.977 |
| 022F | 564 | 550 | 0.975 |
| 031F | 409 | 396 | 0.968 |
| 094F | 62 | 60 | 0.968 |
| 097F | 61 | 59 | 0.967 |
| 095F | 60 | 58 | 0.967 |
| 062F | 146 | 141 | 0.966 |
| 053F | 175 | 169 | 0.966 |
| 047F | 232 | 224 | 0.966 |
| 110F | 52 | 50 | 0.962 |
| 036F | 353 | 337 | 0.955 |
| 044F | 262 | 250 | 0.954 |
| 064F | 135 | 128 | 0.948 |
| 070F | 114 | 108 | 0.947 |
| 068F | 108 | 102 | 0.944 |
| 054F | 176 | 166 | 0.943 |
| 067F | 122 | 115 | 0.943 |
| 106F | 52 | 49 | 0.942 |
| 107F | 52 | 49 | 0.942 |
| 080F | 83 | 78 | 0.940 |
| 088F | 69 | 64 | 0.928 |
| 074F | 107 | 99 | 0.925 |
| 056F | 173 | 160 | 0.925 |
| 045F | 252 | 231 | 0.917 |

**Table S7:**
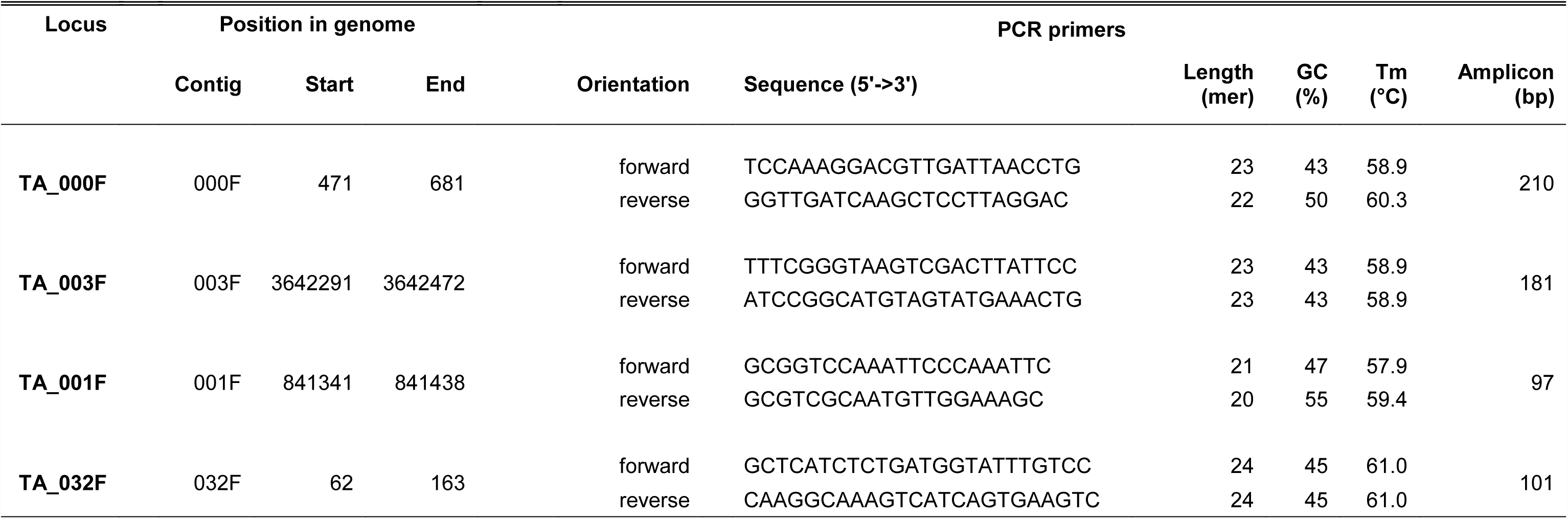
PCR-primers used for verification of selected CNV-loci.

**Table S8:** Data from ddPCR-measurements.

| Clone | Primer | Estimated copies/nl |  |  | Droplet counts |  |  |
| --- | --- | --- | --- | --- | --- | --- | --- |
|  |  | Estimate | ConfMax | ConfMin | Positives | Negatives | Accepted |
| ik1 | TA_001F | 40.1 | 43.6 | 38.3 | 497 | 14340 | 14837 |
| ik1 | TA_003F | 44.1 | 47.8 | 42.1 | 523 | 13708 | 14231 |
| ik1 | TA_032F | 63.4 | 67.9 | 61 | 751 | 13574 | 14325 |
| ik1 | TA_000F | 0.58 | 1.12 | 0.38 | 7 | 14308 | 14315 |
| neg | TA_001F | 0 | 0.22 | 0 | 0 | 15814 | 15814 |
| neg | TA_003F | 0 | 0.24 | 0 | 0 | 14881 | 14881 |
| neg | TA_032F | 0.07 | 0.35 | 0.02 | 1 | 15833 | 15834 |
| neg | TA_000F | 1.2 | 2 | 1 | 16 | 15155 | 15171 |
| ohj104 | TA_001F | 80.1 | 85 | 77.7 | 1072 | 15206 | 16278 |
| ohj104 | TA_003F | 58.2 | 62.6 | 56 | 700 | 13791 | 14491 |
| ohj104 | TA_032F | 0.15 | 0.48 | 0.06 | 2 | 15859 | 15861 |
| ohj104 | TA_000F | 0.53 | 1.04 | 0.35 | 7 | 15476 | 15483 |
| ohj105 | TA_001F | 59.7 | 64 | 57.4 | 715 | 13746 | 14461 |
| ohj105 | TA_003F | 45.5 | 49.8 | 43.4 | 444 | 11256 | 11700 |
| ohj105 | TA_032F | 1.3 | 2 | 1 | 16 | 14976 | 14992 |
| ohj105 | TA_000F | 1.3 | 2 | 1 | 16 | 14668 | 14684 |
| ohj13 | TA_001F | 53.2 | 57.3 | 51.1 | 658 | 14229 | 14887 |
| ohj13 | TA_003F | 48.2 | 52.1 | 46.3 | 602 | 14382 | 14984 |
| ohj13 | TA_032F | 44.9 | 48.6 | 43 | 552 | 14195 | 14747 |
| ohj13 | TA_000F | 0.76 | 1.34 | 0.54 | 10 | 15477 | 15487 |
| ohj22 | TA_001F | 89.6 | 94.8 | 87 | 1160 | 14650 | 15810 |
| ohj22 | TA_003F | 72 | 76.8 | 69.6 | 865 | 13705 | 14570 |
| ohj22 | TA_032F | 0 | 0.23 | 0 | 0 | 15068 | 15068 |
| ohj22 | TA_000F | 0.57 | 1.11 | 0.38 | 7 | 14428 | 14435 |
| ohj7 | TA_001F | 61 | 65.4 | 58.7 | 726 | 13645 | 14371 |
| ohj7 | TA_003F | 50 | 54 | 48 | 608 | 13992 | 14600 |
| ohj7 | TA_032F | 109.9 | 115.5 | 107 | 1463 | 14942 | 16405 |
| ohj7 | TA_000F | 1.27 | 1.97 | 0.98 | 17 | 15772 | 15789 |
| ohj7i3n10 | TA_001F | 68.5 | 73.2 | 66.1 | 815 | 13589 | 14404 |
| ohj7i3n10 | TA_003F | 56.7 | 61.1 | 54.4 | 636 | 12888 | 13524 |
| ohj7i3n10 | TA_032F | 133.4 | 139.9 | 130.1 | 1629 | 13570 | 15199 |
| ohj7i3n10 | TA_000F | 0.59 | 1.1 | 0.4 | 8 | 15982 | 15990 |
| ohj7i3n2 | TA_001F | 41 | 44.5 | 39.2 | 539 | 15200 | 15739 |
| ohj7i3n2 | TA_003F | 38.9 | 42.4 | 37.2 | 502 | 14917 | 15419 |
| ohj7i3n2 | TA_032F | 127.6 | 133.7 | 124.5 | 1680 | 14664 | 16344 |
| ohj7i3n2 | TA_000F | 0.76 | 1.33 | 0.54 | 10 | 15523 | 15533 |
| ohj7i3n5 | TA_001F | 46.2 | 50.1 | 44.2 | 529 | 13218 | 13747 |
| ohj7i3n5 | TA_003F | 40.7 | 44.5 | 38.7 | 430 | 12222 | 12652 |
| ohj7i3n5 | TA_032F | 126.7 | 133.3 | 123.4 | 1429 | 12565 | 13994 |
| ohj7i3n5 | TA_000F | 0.43 | 0.94 | 0.26 | 5 | 13668 | 13673 |
| ohj7i3n7 | TA_001F | 65.1 | 70.9 | 62.1 | 481 | 8458 | 8939 |
| ohj7i3n7 | TA_003F | 46.5 | 50.3 | 44.6 | 600 | 14874 | 15474 |
| ohj7i3n7 | TA_032F | 156 | 163 | 152 | 1919 | 13578 | 15497 |
| ohj7i3n7 | TA_000F | 0.51 | 0.99 | 0.34 | 7 | 16221 | 16228 |
| ohj82 | TA_001F | 74.5 | 79.2 | 72.1 | 949 | 14519 | 15468 |
| ohj82 | TA_003F | 65.1 | 69.8 | 62.7 | 727 | 12783 | 13510 |
| ohj82 | TA_032F | 0.07 | 0.35 | 0.02 | 1 | 16175 | 16176 |
| ohj82 | TA_000F | 1.1 | 1.79 | 0.83 | 14 | 14950 | 14964 |
| ohj96 | TA_001F | 57 | 61.3 | 54.8 | 688 | 13859 | 14547 |
| ohj96 | TA_003F | 45.7 | 49.4 | 43.8 | 590 | 14901 | 15491 |
| ohj96 | TA_032F | 54.2 | 58.3 | 52.1 | 653 | 13858 | 14511 |
| ohj96 | TA_000F | 0.33 | 0.79 | 0.19 | 4 | 14106 | 14110 |
| ohj97 | TA_001F | 62 | 66.1 | 59.9 | 857 | 15846 | 16703 |
| ohj97 | TA_003F | 44.5 | 48 | 42.7 | 620 | 16079 | 16699 |
| ohj97 | TA_032F | 55 | 58.9 | 53 | 749 | 15652 | 16401 |
| ohj97 | TA_000F | 0.3 | 0.7 | 0.17 | 4 | 15757 | 15761 |
| ohj98 | TA_001F | 57.4 | 61.7 | 55.2 | 689 | 13772 | 14461 |
| ohj98 | TA_003F | 51.1 | 55.2 | 49.1 | 604 | 13594 | 14198 |
| ohj98 | TA_032F | 1.16 | 1.86 | 0.89 | 15 | 15193 | 15208 |
| ohj98 | TA_000F | 1.3 | 2 | 1 | 16 | 15009 | 15025 |

**Table S9:**
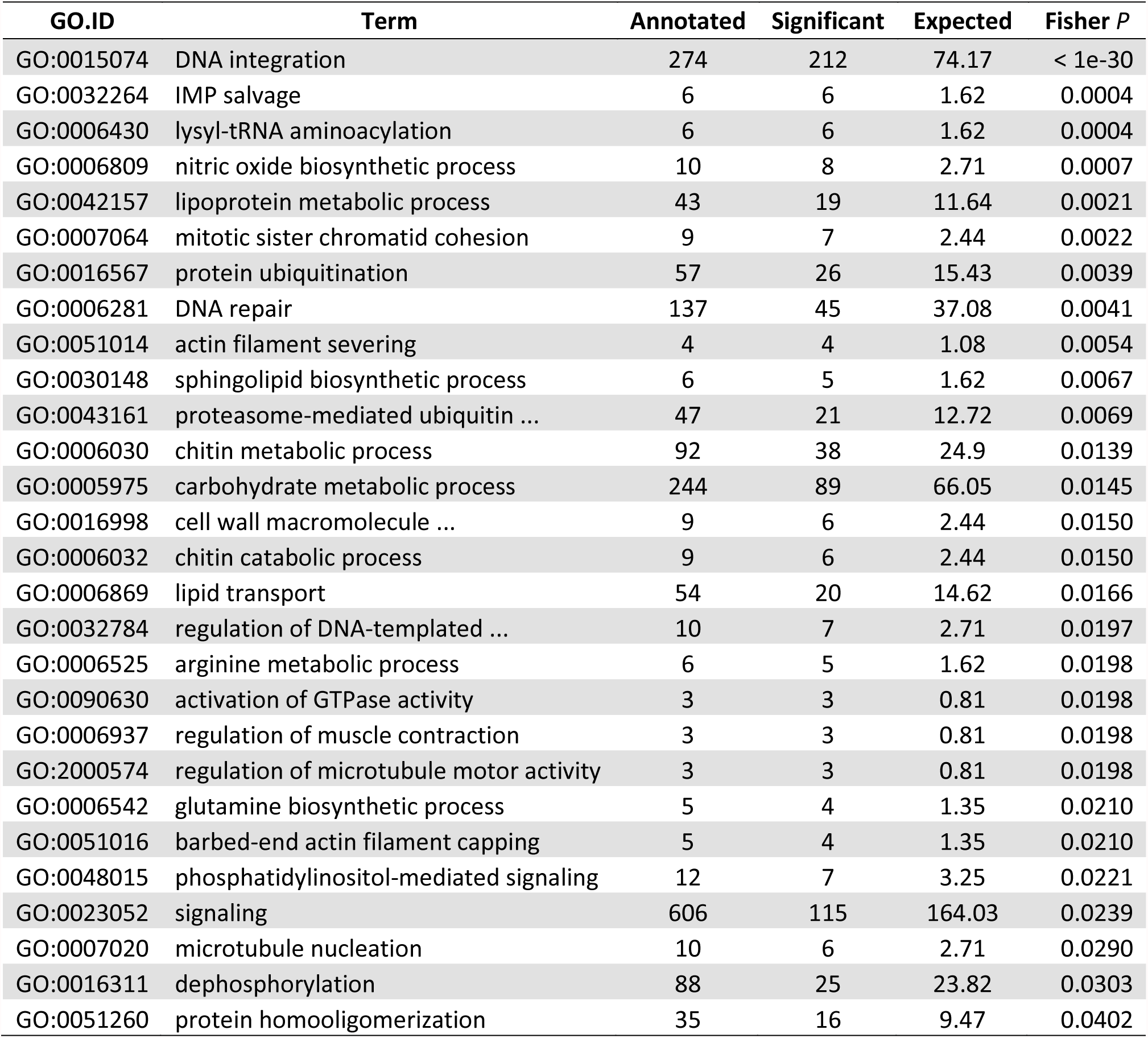
GO enrichment analysis of genes that derived from a duplication event.

| GO.ID | Term | Annotated | Significant | Expected | Fisher *P* |
| --- | --- | --- | --- | --- | --- |
| GO:0015074 | DNA integration | 274 | 212 | 74.17 | < 1e-30 |
| GO:0032264 | IMP salvage | 6 | 6 | 1.62 | 0.0004 |
| GO:0006430 | lysyl-tRNA aminoacylation | 6 | 6 | 1.62 | 0.0004 |
| GO:0006809 | nitric oxide biosynthetic process | 10 | 8 | 2.71 | 0.0007 |
| GO:0042157 | lipoprotein metabolic process | 43 | 19 | 11.64 | 0.0021 |
| GO:0007064 | mitotic sister chromatid cohesion | 9 | 7 | 2.44 | 0.0022 |
| GO:0016567 | protein ubiquitination | 57 | 26 | 15.43 | 0.0039 |
| GO:0006281 | DNA repair | 137 | 45 | 37.08 | 0.0041 |
| GO:0051014 | actin filament severing | 4 | 4 | 1.08 | 0.0054 |
| GO:0030148 | sphingolipid biosynthetic process | 6 | 5 | 1.62 | 0.0067 |
| GO:0043161 | proteasome-mediated ubiquitin ... | 47 | 21 | 12.72 | 0.0069 |
| GO:0006030 | chitin metabolic process | 92 | 38 | 24.9 | 0.0139 |
| GO:0005975 | carbohydrate metabolic process | 244 | 89 | 66.05 | 0.0145 |
| GO:0016998 | cell wall macromolecule ... | 9 | 6 | 2.44 | 0.0150 |
| GO:0006032 | chitin catabolic process | 9 | 6 | 2.44 | 0.0150 |
| GO:0006869 | lipid transport | 54 | 20 | 14.62 | 0.0166 |
| GO:0032784 | regulation of DNA-templated ... | 10 | 7 | 2.71 | 0.0197 |
| GO:0006525 | arginine metabolic process | 6 | 5 | 1.62 | 0.0198 |
| GO:0090630 | activation of GTPase activity | 3 | 3 | 0.81 | 0.0198 |
| GO:0006937 | regulation of muscle contraction | 3 | 3 | 0.81 | 0.0198 |
| GO:2000574 | regulation of microtubule motor activity | 3 | 3 | 0.81 | 0.0198 |
| GO:0006542 | glutamine biosynthetic process | 5 | 4 | 1.35 | 0.0210 |
| GO:0051016 | barbed-end actin filament capping | 5 | 4 | 1.35 | 0.0210 |
| GO:0048015 | phosphatidylinositol-mediated signaling | 12 | 7 | 3.25 | 0.0221 |
| GO:0023052 | signaling | 606 | 115 | 164.03 | 0.0239 |
| GO:0007020 | microtubule nucleation | 10 | 6 | 2.71 | 0.0290 |
| GO:0016311 | dephosphorylation | 88 | 25 | 23.82 | 0.0303 |
| GO:0051260 | protein homooligomerization | 35 | 16 | 9.47 | 0.0402 |

**Table S10:** GO enrichment analysis of genes derived from a duplication event and found within CNV regions.

| GO.ID | Term | Annotated | Significant | Expected | Fisher<br>*P* |
| --- | --- | --- | --- | --- | --- |
| GO:0007165 | signal transduction | 591 | 14 | 9.54 | 0.0007 |
| GO:0048015 | phosphatidylinositol-mediated signaling | 12 | 3 | 0.19 | 0.0008 |
| GO:0046854 | phosphatidylinositol phosphorylation | 14 | 3 | 0.23 | 0.0013 |
| GO:0008285 | negative regulation of cell proliferatio... | 5 | 2 | 0.08 | 0.0025 |
| GO:0042147 | retrograde transport endosome to Golgi | 10 | 2 | 0.16 | 0.0107 |
| GO:0016573 | histone acetylation | 14 | 2 | 0.23 | 0.0207 |
| GO:0005991 | trehalose metabolic process | 20 | 2 | 0.32 | 0.0406 |
| GO:0007017 | microtubule-based process | 191 | 11 | 3.08 | 0.0462 |
| GO:0098535 | de novo centriole assembly ... | 3 | 1 | 0.05 | 0.0477 |
| GO:2000574 | regulation of microtubule motor activity | 3 | 1 | 0.05 | 0.0477 |
| GO:0007040 | lysosome organization | 3 | 1 | 0.05 | 0.0477 |

## Notes

### Competing Interest Statement

The authors have declared no competing interest.

### Summary of Updates

This version additionally contains GO enrichment analysis of duplicated genes. Everything else is the same as in version 1.

